# High Drug-Loading Rucaparib-FdUMP Nanocarriers for Colorectal Cancer Combination Therapy

**DOI:** 10.64898/2026.05.25.727099

**Authors:** Anna Erika Weber, Christian Ritschel, Claus Feldmann, Susanne Kossatz

## Abstract

Combination chemotherapy remains the standard of care for advanced colorectal cancer, yet the clinical benefit is limited by off-target toxicity and insufficient tumor drug exposure. Co-administration of 5-fluorouracil (5-FU) with poly(ADP-ribose) polymerase (PARP) inhibitors such as rucaparib (RUC) can potentiate 5-FU-induced DNA damage by suppressing DNA repair but achieving robust co-delivery at defined ratios and high payload remains challenging. Here, we report high drug-loading core@shell nanocarriers (RUC@[ZrO][FdUMP]) that co-deliver RUC and the active 5-FU metabolite fluoro-2′-deoxyuridine 5′-monophosphate (FdUMP). Nanocarriers were synthesized via a solvent-antisolvent approach to form a tocopherol phosphate (TocP)-stabilized RUC core followed by the growth of an amorphous [ZrO][FdUMP] shell, yielding an overall drug load of 59 wt%. Dual fluorescence (shell incorporation of DUT647 and intrinsic RUC emission) enabled quantitative tracking and revealed rapid uptake in HCT116 and HT29 colorectal cancer cells and tumor accumulation in a subcutaneous HT29 xenograft. In vitro, RUC@[ZrO][FdUMP] showed cytotoxicity comparable to the free-drug combination, induced S-phase arrest, and increased γH2AX foci, consistent with replication stress and DNA double-strand break formation. Drug–drug interaction analysis demonstrated synergistic activity of the RUC/FdUMP pair in colorectal cancer cells. Together, these results establish RUC@[ZrO][FdUMP] as a high-payload nanocarrier platform for co-delivering synergistic PARP inhibitor/fluoropyrimidine combinations, supporting further development and in vivo evaluation of this co-delivery strategy for colorectal cancer therapy.

**TOC Graphics:** 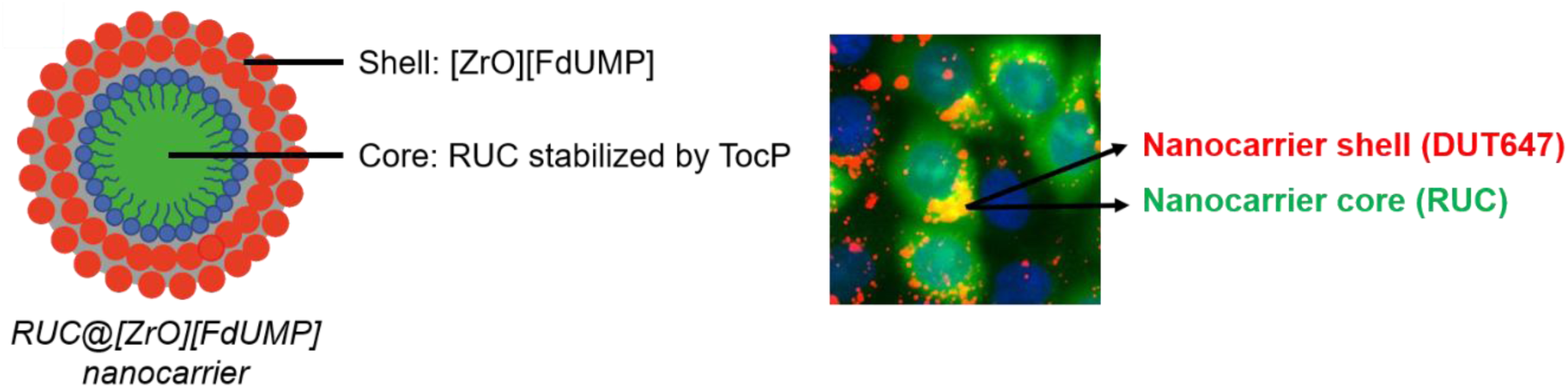

## 1. Introduction

Colorectal cancer is the third most prevalent cancer worldwide, accounting for approximately 900,000 deaths every year [1]. Because symptoms are often nonspecific, diagnosis is delayed, resulting in a poor prognosis with a 5-year survival rate of 15% at stage IV [2]. Treatment of advanced stages usually involves combination chemotherapy, such as FOLFOX (5-fluorouracil (5-FU), leucovorin, and oxaliplatin) or FOLFIRI (5-FU, leucovorin, and irinotecan) [3]. However, the efficacy of these drug cocktails is limited by systemic toxicity and insufficient tumor-selective drug delivery, resulting in severe side effects [4]. Therefore, new treatment strategies are needed in order to improve the efficacy of colorectal cancer therapy.

The anti-metabolite 5-FU plays a pivotal part in the treatment of colorectal cancer and many other solid tumors [5, 6]. It is employed either as a single treatment or as a drug cocktail in combination with other drugs [3]. Once taken up by the cell, 5-FU is converted to the active metabolite fluoro-2′-deoxyuridine 5′-monophosphate (FdUMP), which inhibits thymidylate synthase [5, 6]. As a result, low nucleotide levels lead to replication fork stalling, causing DNA damage [5, 6]. Furthermore, FdUMP gets converted to fluorodeoxyuridine triphosphate (FdUTP), which can be misincorporated into the DNA during synthesis [5, 6]. Consequently, DNA repair pathways such as base excision repair (BER) are activated [5]. Since BER involves the activation of poly(ADP-ribose) polymerase (PARP), the repair of 5-FU/FdUMP-induced DNA damage can be inhibited by PARP inhibitors. Moreover, tumor cells can acquire 5-FU resistance by overexpressing DNA repair enzymes, rendering them vulnerable to PARP inhibition [7].

The four PARP inhibitors olaparib, rucaparib (RUC), niraparib, and talazoparib have been approved as monotherapy for BRCA-mutated cancers, including breast and ovarian cancer [8–11]. PARP inhibitors bind the DNA repair enzyme PARP and induce lethal DNA double-strand breaks (DSB) through PARP trapping [12]. In homologous-repair-deficient tumor cells, the resulting DNA DSB cannot be repaired, leading to cell death [13]. However, recent evidence suggests that PARP inhibitors may also exhibit anti-tumor properties in homologous-repair-proficient tumor cells, when combined with other DNA-damaging agents such as 5-FU [14–16].

Although preclinical studies showed enhanced cytotoxicity of 5-FU if combined with PARP inhibitors, clinical studies are limited [14–16]. One of the first pieces of evidence showed that RUC enhances the DNA-damaging effects of 5-FU in acute leukemic cells, while data from Augustine et al. suggest a greater-than-additive effect in HCT116 colorectal cancer cells [16, 17]. Since about 15 % of colorectal cancer patients have microsatellite instability/mismatch repair deficiency [18], a significant number of colorectal cancer patients might be vulnerable to DNA repair inhibition by PARP inhibitors. Therefore, a phase II trial (NCT00912743) tested olaparib as monotherapy for metastatic colorectal cancer patients with microsatellite instability but did not find a prolonged progression-free or overall survival [19]. It was suggested that the effect be further investigated in combination with other drugs [19]. In 2025, a phase I/II trial (NCT03337087) showed sustained disease response in metastatic gastrointestinal cancer patients after being given a combination of RUC, 5-FU, leucovorin and liposomal irinotecan [20].

5-FU- and PARP inhibitor-based treatment schedules, however, are associated with severe hematologic and gastrointestinal toxicities [6, 21, 22]. Encapsulating the drugs into nanoparticles could be an efficient strategy to reduce off-target binding and increase drug bioavailability [23]. Several strategies for nanoformulations incorporating two active substances have been studied. One such approach is the use of liposomes, which are capable of encapsulating both hydrophilic and lipophilic drugs [24, 25]. Hydrophilic substances are encapsulated in the hydrophilic core of the liposome, while lipophilic compounds are integrated into the lipid bilayer. A notable example is the FDA- and EMA-approved liposomal formulation CPX-351 (brand name VYXEOS®), which combines the chemotherapeutic agents daunorubicin (lipophilic) and cytarabine (hydrophilic) for the treatment of secondary acute myeloid leukemia [26, 27]. A limitation of liposomal formulations is their restricted drug loading capacity. Liposomes typically exhibit a drug-to-lipid weight ratio of less than 0.5, which corresponds to less than 33% of drug by weight of the total nanocarrier mass [28, 29].

To address the loading limitations of liposomal formulations, we are focusing on developing inorganic-organic hybrid nanocarriers with a higher drug content (> 50 wt%). The inorganic component is a metal cation (e.g. [ZrO]^2+^), which binds to an organic drug anion (e.g. gemcitabine monophosphate [GMP]^2-^ or [FdUMP]^2-^). Efficient drug release from these nanocarriers is achieved through hydrolytic cleavage by phosphatases or through acid catalysis [30], both enabled within endolysosomal compartments of cells. Recently, we successfully presented nanocarriers incorporating gemcitabine monophosphate for the treatment of pancreatic cancer and bedaquiline for tuberculosis [31, 32]. A combination of drugs with different chemical properties in a single nanocarrier is still particularly challenging. Nevertheless, we also successfully synthesized a new type of core@shell nanocarrier with the chemotherapeutics irinotecan (ITC) and FdUMP. However, with a drug content of 32 wt%, these nanocarriers exhibit a drug loading comparable to that of established liposomal formulations [33].

Nanomedicine formulations containing PARP inhibitors are a small but emerging field of interest [34–38]. The majority of studies focused on the encapsulation of the PARP inhibitors olaparib and talazoparib [37]. For RUC, only one encapsulation approach in liposomes has been reported [35]. Additionally, the use of carbon nanotubes (CNT) has been suggested for RUC delivery [39]. However, these concepts lack sufficient drug load (< 1% by weight), or have not yet been tested in biological media [35, 39]. For FdUMP, however, nanocarrier encapsulation methods were already examined in more detail by us and other groups [40–42]. In order to establish a nanocarrier-based 5-FU and RUC combination therapy approach, we developed a nanocarrier encapsulating FdUMP, the active metabolite of 5-FU, and the PARP inhibitor RUC using a nanocarrier design that is tailored towards high drug load. The combination of RUC and FdUMP in nanocarriers is completely novel in the literature. Here, we report the synthesis and biological evaluation of RUC-FdUMP nanocarriers with high drug load, aiming to improve the efficacy of colorectal cancer treatment.

## 2. Materials and Methods

A full description of materials and methods can be found in the SI.

## 3. Results

### 3.1. Synthesis of RUC@[ZrO][FdUMP] core@shell nanocarriers

In vivo application of drug nanocarriers requires a drug load as high as possible to limit liquid volume and total number of nanocarriers to achieve clinically relevant drug concentrations. This is even more relevant for nanocarriers containing drug cocktails of two or more drugs. Aiming at drug loads as high as possible, the synthesis of the RUC@[ZrO][FdUMP] nanocarriers was performed using a solvent-antisolvent approach (Figure 1) [43, 44]. As the solubility of RUC as a lipophilic drug in water is very low, we used DMSO as the solvent [45]. Thereafter, this concentrated solution of RUC in DMSO was injected into an excess of cooled water (0 °C) as the anti-solvent. Whereas DMSO and water are miscible in any ratio, RUC is insoluble in water, so that immediate nucleation of RUC nanocarriers occurred after injection. The as-formed RUC nanocarriers are colloidally of limited stability as the interaction between the lipophilic particle surfaces is stronger as the interaction of the RUC nanocarriers with water as a polar solvent. To prevent agglomeration, tocopherol phosphate (TocP, 0.2 eq. of RUC) was added to the anti-solvent prior to RUC nucleation for surface functionalization (Figure 1a). TocP – a derivative of vitamin E – is a biocompatible surfactant adsorbing on the lipophilic RUC nanocarriers with its lipophilic tail and the polar phosphate group directed to the water phase. As a result, a colloidally stable suspension of TocP-stabilized RUC nanocarriers was obtained (Figure 1b) as indicated by a zeta potential of –30 to –70 mV at pH 4-10 (Figure 1d). These TocP-stabilized RUC nanocarriers exhibited a size of 24 ± 8 nm (*SI: Figures S1-S6)*.

**Figure 1.**
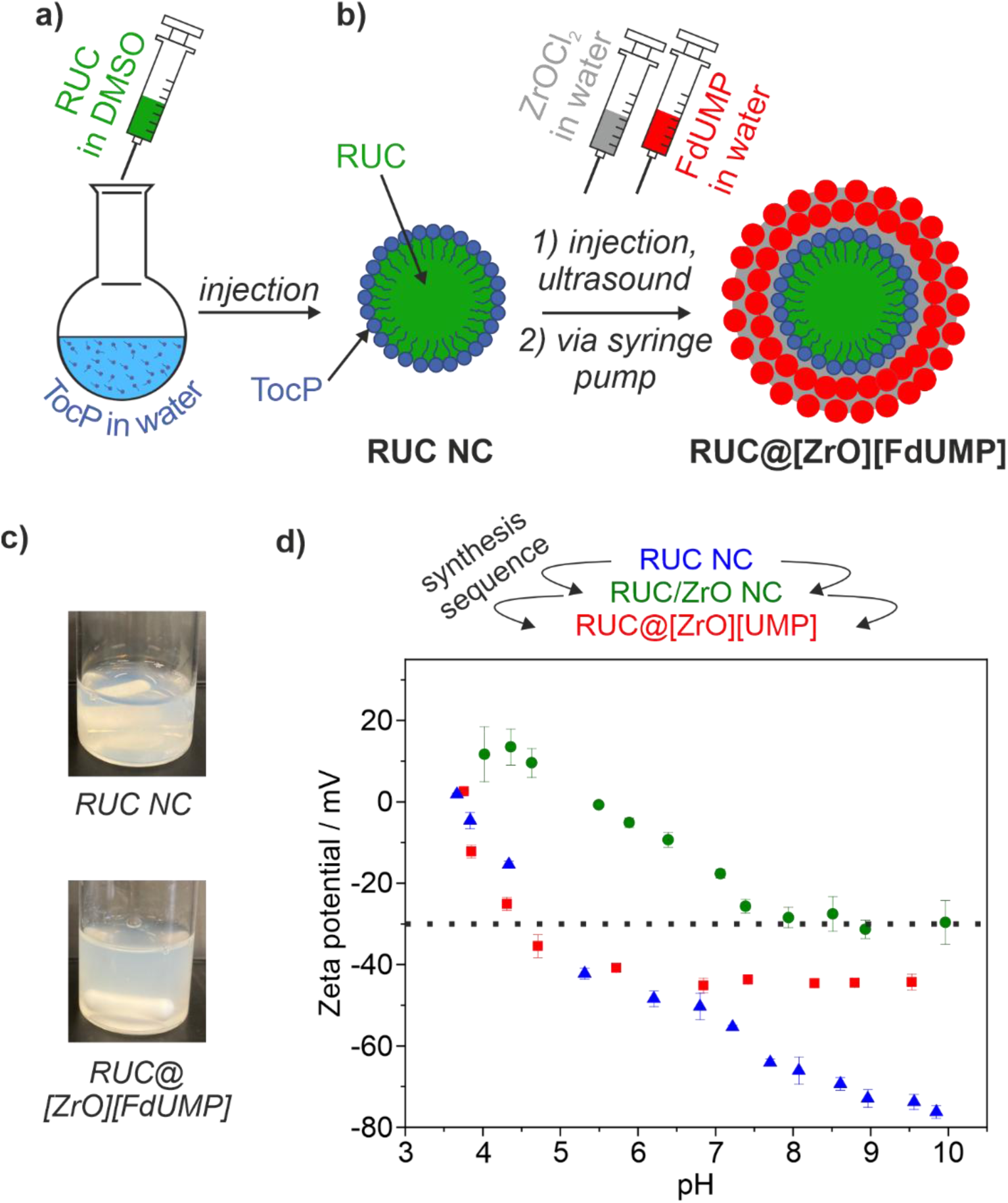
Scheme illustrating the synthesis of RUC@[ZrO][FdUMP] core@shell nanocarriers. **a)** Solvent-antisolvent approach to obtain TocP-stabilized RUC nanocarriers (RUC NC). **b)** Formation of [ZrO][FdUMP] shell around the RUC particle core (RUC: rucaparib; FdUMP: 5-fluoro-2’-deoxyuridine-5’-monophosphate); **c)** Photos of the respective suspensions. **d)** Zeta-potential analysis to follow the course of the reaction (*see SI for more analytical data*).

Since one major drawback of 5-FU is its rapid metabolic degradation upon administration [6], we directly encapsulated its active metabolite, FdUMP, in the nanocarrier. Besides the benefit of directly using the active metabolite, FdUMP also fits well with our concepts to realize nanocarriers [30,32,33]. To establish the particle shell, aqueous solutions of ZrOCl_2_×8H_2_O and Na_2_(FdUMP) were added (Figure 1c). First of all, ZrOCl_2_×8H_2_O was added to initiate a binding of zirconyl cations ([ZrO]^2+^) to the phosphate groups of surface-bound TocP (*see SI*). The resulting cation termination reduced the zeta potential to ±0 to –30 mV at pH 5-10 and intermediately also reduced the colloidal stability of the suspension (Figure 1b,d). Thereafter, Na_2_(FdUMP) was added. Having established a thin particle shell, ZrOCl_2_×8H_2_O and Na_2_(FdUMP) solutions were slowly added simultaneously using two different syringe pumps to increase the thickness of the particle shell. With the formation of the [ZrO][FdUMP] particle shell, an anion termination of the nanocarriers was re-established with an increase of the negative surface charging and a zeta potential of –30 to –50 mV at pH 4-10 (Figure 1c,d) [42]. The as-prepared RUC@[ZrO][FdUMP] core@shell nanocarriers were purified by centrifugation/redispersion in/from water to remove excess starting material and salts. Finally, they were dried to powder samples (vacuum, 50 °C) or redispersed in water with sodium citrate (17 mM) to increase the colloidal stability of the suspensions to 4-8 weeks.

### 3.2. Characterization of RUC@[ZrO][FdUMP] core@shell nanocarriers

Due to the high costs of Na_2_(FdUMP) (5 mg ∼ 230 €), the material characterization of the RUC@[ZrO][FdUMP] core@shell nanocarriers was partly performed with Na_2_(UMP) (UMP: uridine monophosphate, 5 g ∼230 €) to result in RUC@[ZrO][UMP] core@shell nanocarriers. The structure of UMP and FdUMP is identical, in principle, except for the absence (UMP) or presence (FdUMP) of the fluorine atoms. Although essential for the anti-tumor activity, the particle size and chemical composition of the RUC@[ZrO][UMP] and RUC@[ZrO][FdUMP] nanocarriers are expected to be similar for given the closely related structure. Size and size distribution of the RUC@[ZrO][UMP] core@shell nanocarriers were examined by dynamic light scattering (DLS) and scanning electron microscopy (SEM) (Figure 2). In this regard, DLS showed a mean hydrodynamic diameter of 126 ± 50 nm. SEM showed nanocarriers with spherical shape and a size range of 30-50 nm (Figure 2a,b). A statistical evaluation of 300 nanocarriers on SEM images resulted in mean diameter of 43 ± 8 nm (Figure 2c).

**Figure 2.**
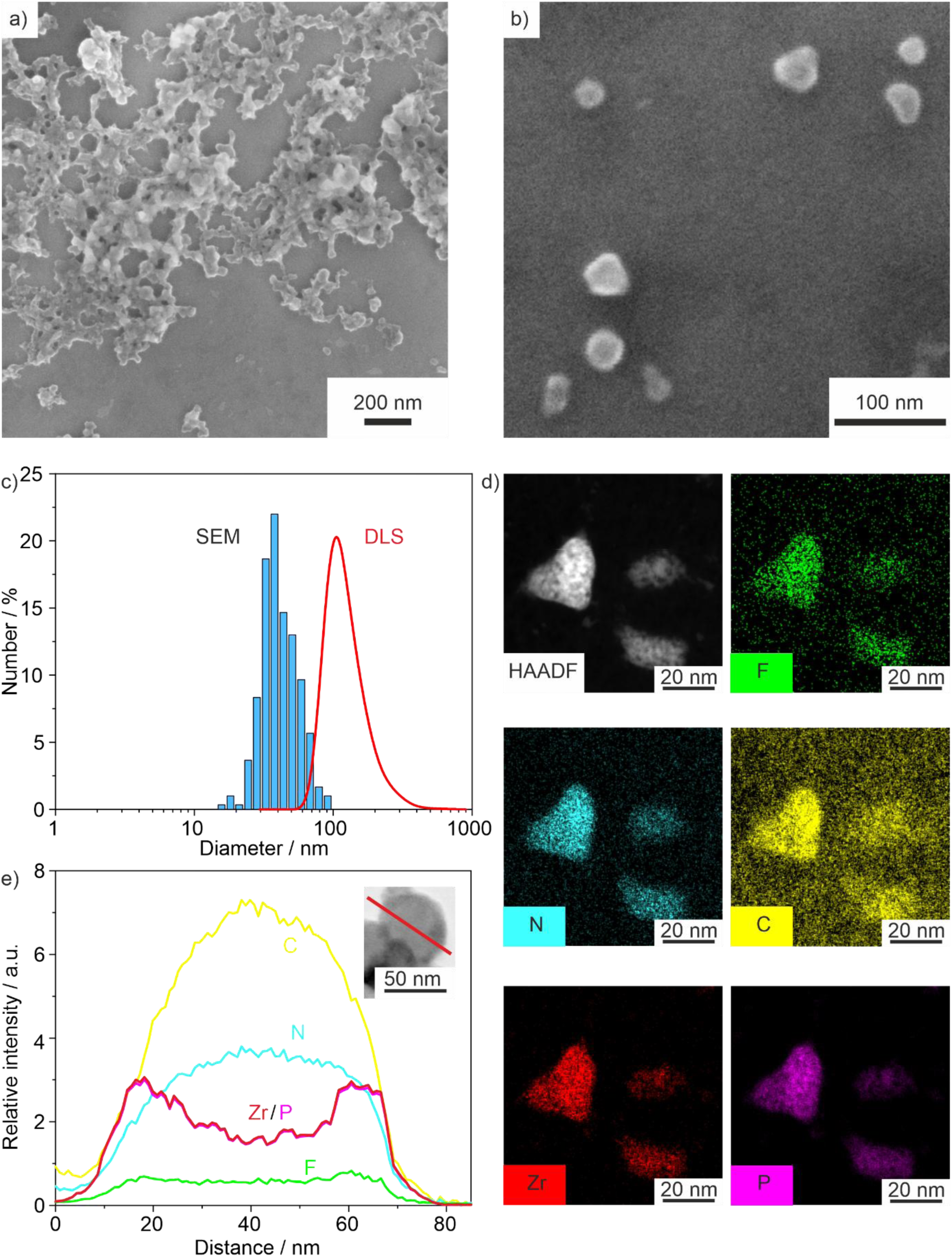
Size and size distribution of RUC@[ZrO][(Fd)UMP] core@shell nanocarriers: **a)** SEM overview image. **b)** SEM detailed image. **c)** Size distribution according to DLS (aqueous suspension) and statistical evaluation of SEM images (300 nanocarriers). **d)** EDXS element maps with HAADF-STEM overview image and elemental distribution of Zr, P, N, C, F. **e)** EDXS linescan (along orange arrow on HAADF-TEM image) with elemental distribution of Zr, P, C, N, F (*see SI for more analytical data*).

Size and structure of the RUC@[ZrO][UMP] core@shell nanocarriers were examined by high-angle annular dark-field scanning transmission electron microscopy (HAADF-STEM) and energy-dispersive X-ray spectroscopy (EDXS) (Figure 2d,e). HAADF-STEM images already indicated the presence of the shell, based on the higher density and electron absorption of ZrO(FdUMP) (Figure 2d). EDXS linescans exhibited a characteristic U-type concentration profile for Zr and P, indicating the presence of these elements only in the particle shell (Figure 2e). Moreover, the elemental maps showed a homogeneous distribution of C, N, F (Figure 2d,e), but again a higher concentration of Zr and P in the particle shell (Figure 2d,e). Similar to SEM and DLS, the TocP-stabilized RUC-core exhibited a diameter of about 25 nm and the [ZrO][FdUMP] shell a diameter of about 18 nm, resulting in a total diameter of about 61 nm.

Finally, the chemical composition of the RUC@[ZrO][UMP] core@shell nanocarriers was analyzed by X-ray powder diffraction (XRD), Fourier-transform infrared (FT-IR) spectroscopy, total organics combustion with thermogravimetry (TG), elemental analysis (EA) and photometry (*SI: Figures S7-S15*). According to XRD, the nanocarriers are amorphous (*SI: Figures S3,S10*), which can be expected for such material and room-temperature synthesis [24–29, 31–33]. Non-crystallinity, on the other hand, is also often beneficial due to a faster drug release in comparison to crystalline materials [35, 39]. FT-IR spectra showed characteristic vibrations that can be attributed to RUC, TocP, UMP and, thus, prove their presence qualitatively (*SI: Figures S3,S10*). Total organics combustion with TG and EA allowed to quantify the composition and drug load of the nanocarriers (*SI: Figures S4,S7,S11*). Thus, EA with C/H/N/S analysis resulted in 32.3 wt% C, 3.1 wt% H, 6.0 wt% N, and 0.0 wt% S (after correction for adsorbed water and DMSO; *SI: Tables S1-S3*). These data were in accordance with a composition RUC@[ZrO][UMP] and calculated values of 32.3 wt% C, 3.1 wt% H, 6.0 wt% N, and 0.0 wt% S. Finally, photometry was applied and resulted in an overall drug load of 59 wt% with an UMP: RUC molar ratio of 1.2: 1 (*SI: Figures S6,S8,S12*). The good agreement of all these analytical data, including total organics combustion with TG, EA and photometry, in sum, reliably confirmed the composition and drug load of the RUC@[ZrO][UMP] core@shell nanocarriers.

To allow fluorescence tracking of the as-prepared RUC@[ZrO][UMP] core@shell nanocarriers with optical imaging, first of all, the intrinsic blue emission of RUC can be used [45]. Moreover, DY 647P1-dUTP (DUT647) – a fluorescent dye functionalized with a uridine triphosphate group – can be used for fluorescence labeling of the particle shell and was added with tiny amounts (40 nmol) to UMP/FdUMP in the synthesis (*SI: Figure S14*). The resulting slightly bluish nanocarriers (due to presence of DUT647), in sum, exhibited both blue emission (RUC-based: *λ_em_(max)* = 483 nm) and red emission (DUT647-based: *λ_em_(max)* = 681 nm) upon blue-light excitation (*λ_ex_(max)* = 360 nm). Notably, the DUT647-based red emission can also be excited with orange-red light (*λ_ex_(max)*= 653 nm; *SI: Figure S14*).

Finally, the stability of RUC@[ZrO][UMP] was evaluated at 37 °C in phosphor-buffered saline (PBS), cell culture medium and cell culture medium with 10% fetal bovine serum (FBS) and penicillin/streptomycin (P/S), to mimic the conditions during cell experiments, with 100 µg/mL of the nanocarriers (Figure 3). Note that these measurements quantify release of the core drug RUC; release of the shell nucleotide (UMP/FdUMP) was not assessed here.

**Figure 3.**
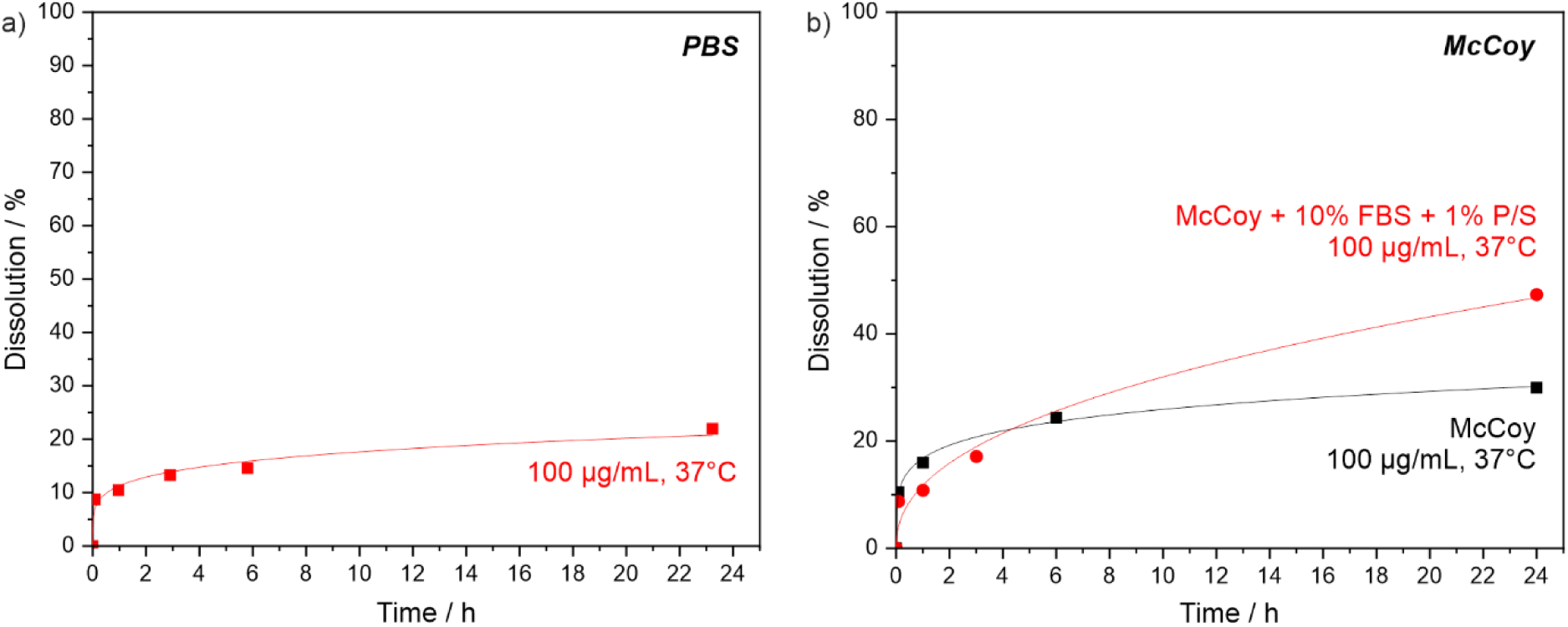
Release of RUC from RUC@[ZrO][(TocP)(UMP)] core@shell nanocarriers in different media. **a)** Release of RUC from 100 µg/mL RUC@[ZrO][(TocP)(UMP)] in phosphor-buffered saline (PBS) at 37 °C. **b)** Release of RUC from 100 µg/mL RUC@[ZrO][(TocP)(UMP)] in pure cell culture medium (McCoýs 5A (modified)) and cell culture medium with 10% fetal bovine serum (FBS) and 1% penicillin/streptomycin (P/S) at 37 °C.

After a rapid release of RUC at the beginning (< 0.5 h), a continuous, slow release of RUC was generally observed. In cell culture medium with 10% FBS and 1% P/S, 10.8 % RUC was released after 1 h, which increased to 17.1 % after 3 h and 47.3 % after 24 h (Figure 3b). However, it should be noted that the drug release from nanocarriers in cells and animals with active metabolism can be substantially different from that in a pure medium. After fast uptake of nanocarriers in cells, the drug release will here predominantly occur inside the respective cells.

### 3.3. Dual fluorescence tracking reveals rapid cellular uptake of RUC@[ZrO][FdUMP/DUT647]

The uptake of nanocarriers by tumor cells is crucial for their therapeutic efficiency. Therefore, we first investigated the cellular uptake of RUC@[ZrO][FdUMP/DUT647] in colon carcinoma cells by tracking the localization of the shell dye DUT647 and the core drug RUC with fluorescence microscopy (Figure 4a). HT29 cells were treated with RUC@[ZrO][FdUMP/DUT647] or equimolar drug concentrations of free RUC and FdUMP for 0.5 h, 4 h and 24 h (Figure 4a). We added FdUMP to the free RUC group to expose the cells to the same drug cocktail as with the nanocarriers. After 0.5 h, signals from DUT647 and RUC were detected in HT29 cells, indicating rapid cellular uptake of RUC@[ZrO][FdUMP/DUT647]. While the signal of the shell dye DUT647 was clustered at the cell membrane and in the cytoplasm, RUC was found in the cytoplasm as well as the nucleus and the nucleoli, similar to our observation for free RUC and FdUMP (Figure 4a). Limited co-localization of DUT647 and RUC indicates that drug release was already initiated within the first 0.5 h of treatment. It should be noted that RUC@[ZrO][FdUMP] showed 10.8 % RUC release after 1 h in complete cell culture medium, which increased to 17.1 % free RUC after 3 h and 47.3 % free RUC after 24 h (Figure 3b). Treatment with free RUC/FdUMP showed a homogeneous cytoplasmic and nuclear RUC signal across all cells and an increasing signal from 0.5 h to 4 h, while no further signal increase was observed after 24 h. For RUC@[ZrO][FdUMP/DUT647], we observed increasing amounts of the shell dye DUT647 from 4 h to 24 h, which was internalized and clustered in HT29 cells. In parallel, we observed an increasing vesicle-like RUC signal at and close to the nuclear membrane, starting at 0.5 h, but becoming more pronounced at 4 h and 24 h. This punctuated RUC signal was largely co-localized with the DUT647 signal, suggesting accumulation of intact RUC@[ZrO][FdUMP] in vesicular structures over time, representing a drug sequestration for later release.

**Figure 4.**
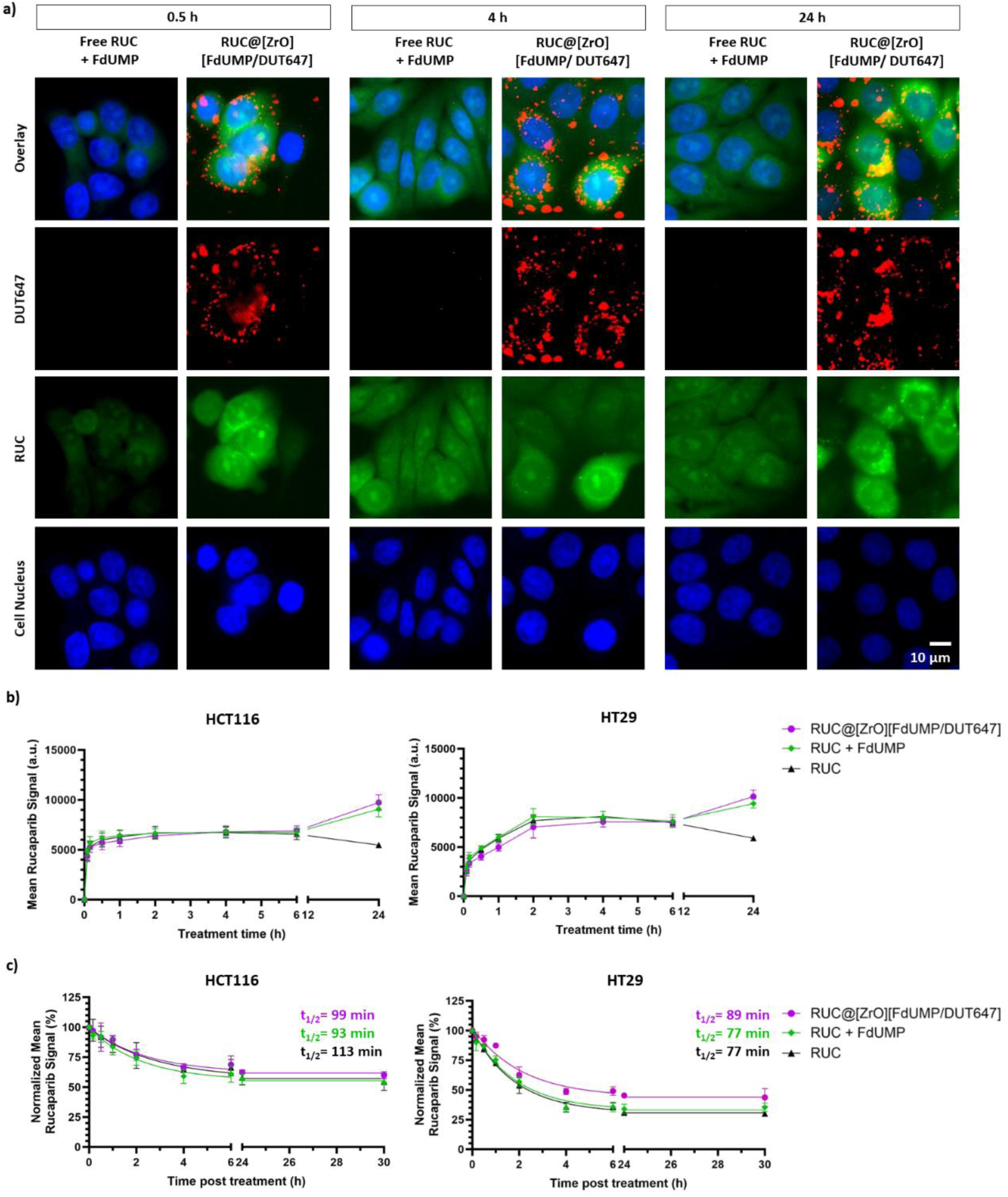
Uptake of RUC@[ZrO][FdUMP/DUT647] in colon carcinoma cells. **a)** Representative microscopy images of HT29 cells treated with 5.65 µg/mL RUC@[ZrO][FdUMP/DUT647] (containing 5 µM RUC and 5.7 µM FdUMP) or 5 µM free RUC and 5.7 free FdUMP as control after 0.5 h, 4 h and 24 h treatment. The shell dye DUT647 of RUC@[ZrO][FdUMP/DUT647] (red), RUC (green) and cell nuclei (blue) are displayed. **b)** Mean fluorescence intensity (MFI) of intracellular RUC after treatment with 0.2 µg/mL RUC@[ZrO][FdUMP/DUT647] (containing 0.2 µM RUC and 0.2 µM FdUMP), 0.2 µM free RUC and 0.2 µM free FdUMP or 0.2 µM free RUC measured up to 24 h. **c)** Intracellular retention of RUC as shown by normalized mean fluorescence intensity of RUC after 4 h treatment with 0.2 µg/mL RUC@[ZrO][FdUMP/DUT647] (containing 0.2 µM RUC and 0.2 µM FdUMP), 0.2 µM free RUC and 0.2 µM free FdUMP or 0.2 µM free RUC as controls and subsequent post-incubation in fresh medium. Curve fitting and half-life calculations were done in GraphPad Prism using a one-phase decay least-squares regression. Values represent mean ± SD of three biological replicates.

To determine the kinetics of RUC@[ZrO][FdUMP/DUT647] uptake, intracellular RUC levels were further quantified using flow cytometry. Therefore, HCT116 and HT29 cells were treated with either RUC@[ZrO][FdUMP/DUT647], equimolar concentrations of free RUC and free FdUMP or free RUC alone for up to 24 h (Figure 4b). Already 5 min after RUC@[ZrO][FdUMP/DUT647] treatment, a RUC signal could be detected in HCT116 and HT29 cells (Figure 4b). The intracellular RUC signal further increased with longer incubation times and reached a plateau between 2 h and 6 h. RUC@[ZrO][FdUMP/DUT647] treatment had similar RUC uptake kinetics compared to the free combination and single drug treatment up to 6 h, despite the vast difference in size and architecture of a nanocarrier versus small molecule delivery. After 24 h treatment, RUC@[ZrO][FdUMP/DUT647] and the combination of free drugs showed a further, comparable increase in RUC uptake. Single treatment with free RUC, on the other hand, showed a decrease in RUC signal from 6 h to 24 h of treatment. Moreover, cells were larger and more granular after 24 h treatment with RUC and FdUMP, but no change in morphology was observed after RUC single treatment (*SI: Figure S15*). Hence, the increased RUC uptake after 24 h of RUC@[ZrO][FdUMP/DUT647] and free-drug combination treatment can be attributed to the effect of FdUMP. In other words, FdUMP from the nanocarrier shell enhanced the uptake of RUC coming from the nanocarrier core, highlighting the beneficial interactions of the two drugs. The reduced uptake after 24 h RUC treatment could be explained by the metabolization of RUC inside the cell.

Similarly, we compared the retention of RUC after RUC@[ZrO][FdUMP/DUT647] or free-drug treatment in tumor cells (Figure 4c). HCT116 and HT29 cells were treated with either RUC@[ZrO][FdUMP/DUT647], equimolar concentrations of free RUC and FdUMP or RUC alone as controls for 4 h. Afterwards, cells were incubated in fresh medium up to 30 h post-treatment, and the intracellular RUC signal was measured using flow cytometry to study drug retention. The intracellular RUC level in HCT116 and HT29 cells was exponentially decreasing and reached a plateau 4 h post-treatment (Figure 4c). In HCT116 cells, the retention of RUC was comparable between RUC@[ZrO][FdUMP/DUT647] treatment and the combination treatment of free RUC and FdUMP, as well as the single drug treatment with RUC. The intracellular half-life of RUC, until the signal reached a plateau, was 99 min after RUC@[ZrO][FdUMP/DUT647] treatment, 93 min after free combination treatment with RUC and FdUMP, and 113 min after free single drug treatment with RUC. In comparison, HT29 cells exhibited lower RUC levels after free-drug treatments than HCT116 cells. Higher levels of RUC were retained after RUC@[ZrO][FdUMP/DUT647] treatment compared to free RUC treatments in HT29 cells. Consequently, the intracellular half-life of RUC, until the signal reached a plateau, in HT29 cells was 89 min after RUC@[ZrO][FdUMP/DUT647] treatment, 77 min after free combination treatment with RUC and FdUMP, and 77 min after free single drug treatment with RUC.

### 3.4. Comparable cytotoxicity and synergistic potential of RUC@[ZrO][FdUMP/DUT647] and the free drugs RUC and FdUMP

Before evaluating the effect of drug-loaded nanocarriers, we tested the cytotoxic effects of the free drugs RUC and FdUMP. Therefore, the viability of HCT116 and HT29 colon carcinoma cells was measured after 72 h treatment with varying concentrations of RUC, FdUMP or a combination of both drugs using an AlamarBlue HS assay (Figure 5). Solvent solutions were shown to be not cytotoxic (*SI: Figure S16*). In HCT116 and HT29 cells, FdUMP was more cytotoxic than RUC, with an EC_50_ of 20.3 ± 3.8 µM in HCT116 cells and 44.8 ± 12.0 µM in HT29 cells for RUC and an EC_50_ of 2.4 ± 0.6 µM in HCT116 cells and 1.9 ± 0.9 µM in HT29 cells for FdUMP (Figure 5a, Table 1). HCT116 cells were more sensitive to the cytotoxic effect of RUC, FdUMP, and their combination than HT29 cells (Figure 5a). In addition, combination treatments of RUC and FdUMP were tested at different molar ratios of the two drugs in order to identify the optimal drug ratio for the strongest cytotoxic effects. Following treatment with an equimolar ratio of RUC and FdUMP (1:1), HCT116 and HT29 cells exhibited comparable levels of cytotoxicity relative to treatments with either a 2:1 or a 1:2 molar ratio (RUC:FdUMP) (Figure 5a).

**Figure 5.**
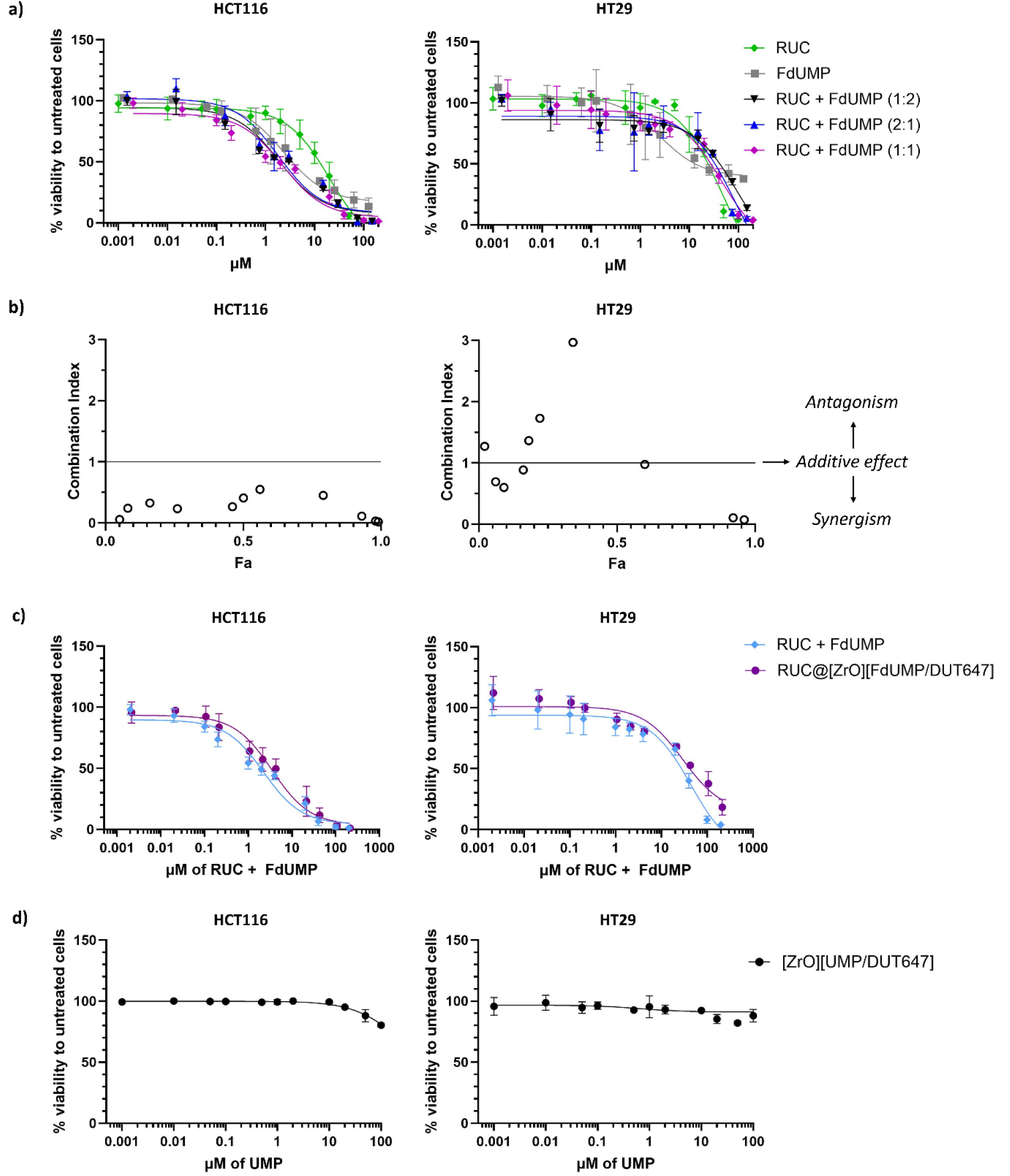
Cytotoxicity of RUC@[ZrO][FdUMP/DUT647] and the free drugs RUC and FdUMP in HCT116 and HT29 colon carcinoma cells. **a)** Cell viability of HCT116 and HT29 cells after 72 h treatment with increasing concentrations of RUC, FdUMP or a combination of both in molar ratios of 1:2, 2:1 or 1:1 (RUC:FdUMP) as measured with an AlamarBlue HS assay. The x-axis shows the total drug concentration in the treatment solution. **b)** Combination index (CI) showing a synergistic (CI < 1), additive (CI = 1) or antagonistic (CI > 1) effect of RUC and FdUMP after equimolar treatment (1:1) based on the cell viability data in (a) and calculated using the software CompuSyn (Fa: inhibitory effect). **c)** Cell viability after 72 h treatment with varying concentrations of RUC@[ZrO][FdUMP/DUT647] (RUC:FdUMP molar ratio 1.00:1.14) or the free-drug combination of RUC and FdUMP (molar ratio 1:1) as measured with an AlamarBlue HS assay. **d)** Cell viability after 72 h treatment with drug-free [ZrO][UMP/DUT647] nanocarriers as measured with an AlamarBlue HS assay. Values represent mean ± SD of three biological replicates. Curve fittings were done in GraphPad Prism using a nonlinear regression model (Dose-response – Inhibition; [Inhibitor] vs. response (three parameters)).

**Table 1.**
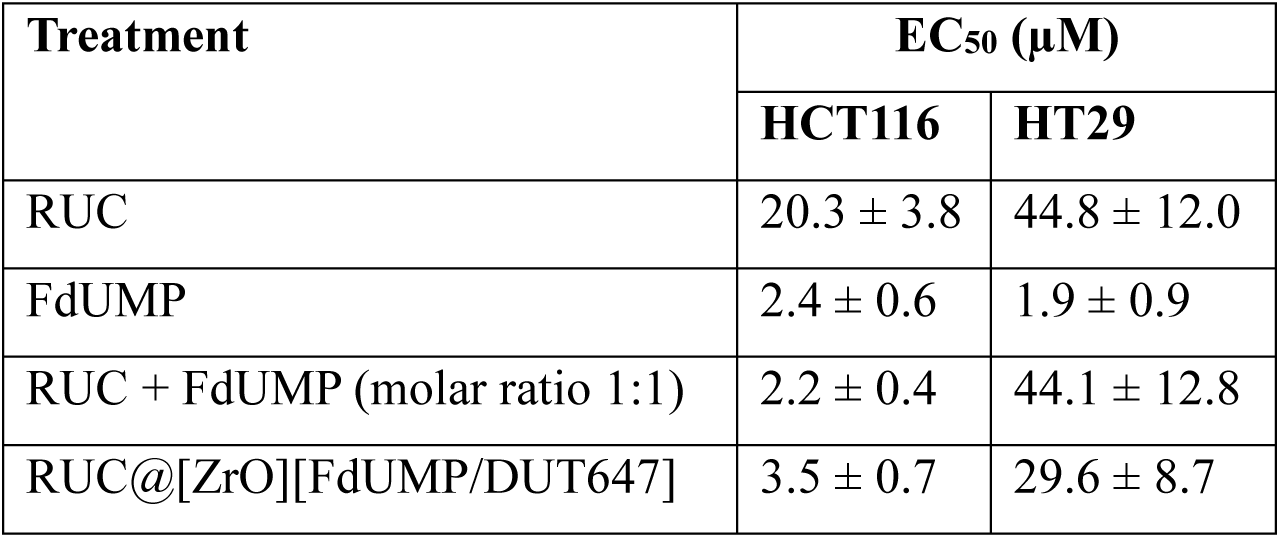
EC_50_ values (µM) with standard errors of the mean after 72 h treatment of RUC@[ZrO][FdUMP/DUT647] nanocarriers or free drugs in HCT116 and HT29 colon carcinoma cells. EC_50_ values were calculated in GraphPad Prism using a nonlinear regression model (Dose-response – Inhibition; [Inhibitor] vs. response (three parameters)).

Based on the cell viability measured after treatment with an equimolar ratio of RUC and FdUMP (1:1) (Figure 5a), the software CompuSyn was used to calculate a combination index (CI) score indicating if the interaction between RUC and FdUMP is synergistic (CI < 1), additive (CI = 1) or antagonistic (CI > 1) (Figure 5b). In HCT116 cells, the combined cytotoxic effect of RUC and FdUMP was found to be synergistic over the entire concentration range (Figure 5b). In HT29 cells, five concentrations were found to be synergistic, one concentration showed an additive effect and four concentrations were antagonistic (Figure 5b). The antagonistic concentration range was primarily between 2 µM and 20 µM, where FdUMP alone was shown to be more cytotoxic than in combination with RUC (Figure 5b). Nevertheless, most concentrations were synergistic or additive in HT29 cells (Figure 5b).

Furthermore, the cytotoxicity of RUC@[ZrO][FdUMP/DUT647] was tested and compared to the free-drug combination of RUC and FdUMP after 72 h (Figure 5c). RUC@[ZrO][FdUMP/DUT647] treatment resulted in similar cytotoxicity as the free-drug combination treatment with free RUC and FdUMP (Figure 5c). EC_50_ values in HCT116 cells were 3.5 ± 0.7 µM for RUC@[ZrO][FdUMP/DUT647] treatment and 2.2 ± 0.4 µM for free RUC and FdUMP treatment, and in HT29 cells 29.6 ± 8.7 µM and 44.1 ± 12.8 µM, respectively (Table 1). In HT29 cells, the EC_50_ value of free RUC and FdUMP treatment was higher than the EC_50_ value of FdUMP single treatment because RUC and FdUMP acted synergistically only at lower and higher concentrations. Drug-free [ZrO][UMP/DUT647] nanocarriers did not notably decrease the viability of HCT116 and HT29 cells (Figure 5d).

### 3.5. RUC@[ZrO][FdUMP/DUT647] treatment arrests cells in S-phase and induces DNA double strand breaks

Next, we investigated the effects of RUC@[ZrO][FdUMP/DUT647] treatment on cell cycle and DNA damage and compared the results to the free-drug treatments with RUC and FdUMP (Figure 6). HCT116 and HT29 cells were treated with 0.1 µg/mL RUC@[ZrO][FdUMP/DUT647], containing 0.1 µM RUC and 0.1 µM FdUMP, or with either 0.1 µM RUC, 0.1 µM FdUMP or a combination of 0.1 µM RUC and 0.1 µM FdUMP for 24 h (Figure 6a, b). RUC treatment alone resulted in a smaller proportion of HT29 cells in the S-phase, while no change in cell cycle distribution was observed for HCT116 cells (Figure 6a, b). On the other hand, FdUMP treatment alone caused G1-phase arrest in HCT116 cells and S-phase arrest in HT29 cells, while reducing the amount of cells reaching the G2-phase. Combination treatment of RUC and FdUMP and RUC@[ZrO][FdUMP] treatment similarily induced S-phase arrest in both cell lines. Since FdUMP single treatment induced S-phase arrest in HT29 cells and rucaparib had the opposite effect, the S-phase arrest after RUC@[ZrO][FdUMP/DUT647] treatment can be mainly attributed to the effect of the shell drug FdUMP. Hence, after RUC@[ZrO][FdUMP/DUT647] treatment, most cells were first arrested in the S-phase and did not reach the G2-phase (Figure 6a, b).

**Figure 6.**
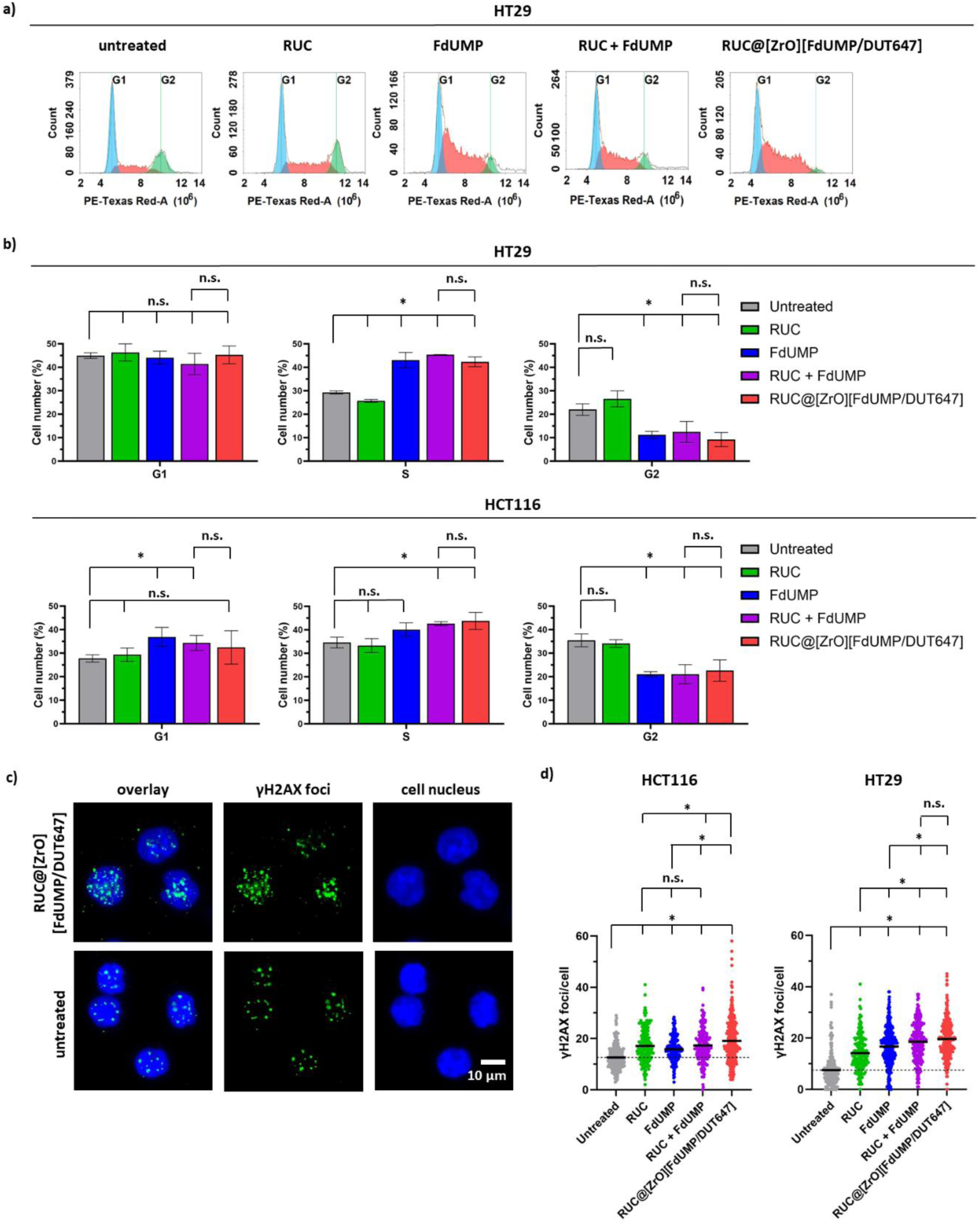
Cell cycle distribution and γH2AX formation after RUC@[ZrO][FdUMP/DUT647] or free RUC and FdUMP treatment in HCT116 and HT29 cells. **a-b)** Flow cytometric cell cycle analysis with propidium iodide stain after treatment of 0.1 µg/mL RUC@[ZrO][FdUMP/DUT647] (containing 0.1 µM RUC and 0.1 µM FdUMP) or free-drug treatments with 0.1 µM RUC, 0.1 µM FdUMP or 0.1 µM RUC and 0.1 µM FdUMP for 24 h or untreated cells. Representative histograms of HT29 cells in a) showing the cell count in the G1- (blue), S-(red), and G2- (green) phase based on the propidium iodide stain intensity of DNA content. Bar graphs in b) showing the percentage of cells in the G1-, S-, and G2-phase after each treatment in HT29 and HCT116 cells. Mean values ± SD of three biological replicates are shown. *p < 0.05; unpaired student’s t-test. **c)** Representative microscopy images of HCT116 cells showing the formation of γH2AX foci (green) after 24 h treatment of 5.65 µg/mL RUC@[ZrO][FdUMP/DUT647] or untreated. The cell nucleus (blue) was counterstained. **d)** Quantification of γH2AX foci in HCT116 and HT29 cells after 24 h treatment of 5.65 µg/mL RUC@[ZrO][FdUMP/DUT647] (containing 5 µM RUC and 5.7 µM FdUMP) or free-drug controls (5 µM RUC, 5 µM FdUMP or 5 µM RUC and 5 µM FdUMP) compared to untreated cells. Scatter plots show the mean of at least 300 cells. *p < 0.05; Mann-Whitney-U test.

Furthermore, the formation of DNA double-strand breaks (DSB) was assessed through immunofluorescent staining of γH2AX foci (Figure 6c, d). All treatments with RUC and/or FdUMP increased the amount of DSB in HCT116 and HT29 cells (Figure 6d). Despite the higher cytotoxicity of FdUMP compared to RUC, RUC single treatment showed the highest increase in γH2AX foci in both cell lines compared to untreated cells (Figure 6d). Thus, RUC treatment was more efficient in DNA DSB formation than FdUMP treatment. RUC@[ZrO][FdUMP/DUT647] treatment induced the highest number of γH2AX foci in both cell lines (Figure 6d). In HCT116 cells, RUC@[ZrO][FdUMP/DUT647] treatment resulted in significantly more DNA DSB than equimolar treatment with the free-drug combination, while no significant difference was observed in HT29 cells (Figure 6d). Hence, RUC@[ZrO][FdUMP/DUT647] treatment is at least as effective as the free-drug combination of RUC and FdUMP in inducing DNA DSB. Taken together, the predominant effect on DNA DSB formation after RUC@[ZrO][FdUMP/DUT647] can be primarily attributed to the effect of RUC from the nanocarrier core.

### 3.6. Tumor uptake of dual fluorescently labeled RUC@[ZrO][FdUMP/DUT647] in an HT29 xenograft

We performed a preliminary in vivo study, to evaluate the uptake of RUC@[ZrO][FdUMP/DUT647] nanocarriers in a HT29 xenograft model. One athymic nude mouse with a subcutaneous HT29 colon carcinoma was intravenously injected with 714 µg RUC@[ZrO][FdUMP/DUT647] in 0.9% NaCl (Figure 7). Fluorescent microscopy of cryosections ex vivo showed a heterogenous tumor uptake of RUC@[ZrO][FdUMP/DUT647] 24 h post-injection (Figure 7a). The nanocarrier shell-dye DUT647 and RUC from the nanocarrier core were found at various locations within the tumor, with the highest signals detected at the tumor rim (Figure 7a). The co-localization of DUT647 and RUC indicated accumulation of labeled nanocarriers, while the distinct staining patterns of DUT647 and RUC suggested successful drug release from the nanocarrier in vivo (Figure 7b). Microscopy of a tumor without RUC@[ZrO][FdUMP/DUT647] injection confirmed the specificity of both signals (*SI: Figure S17*).

**Figure 7.**
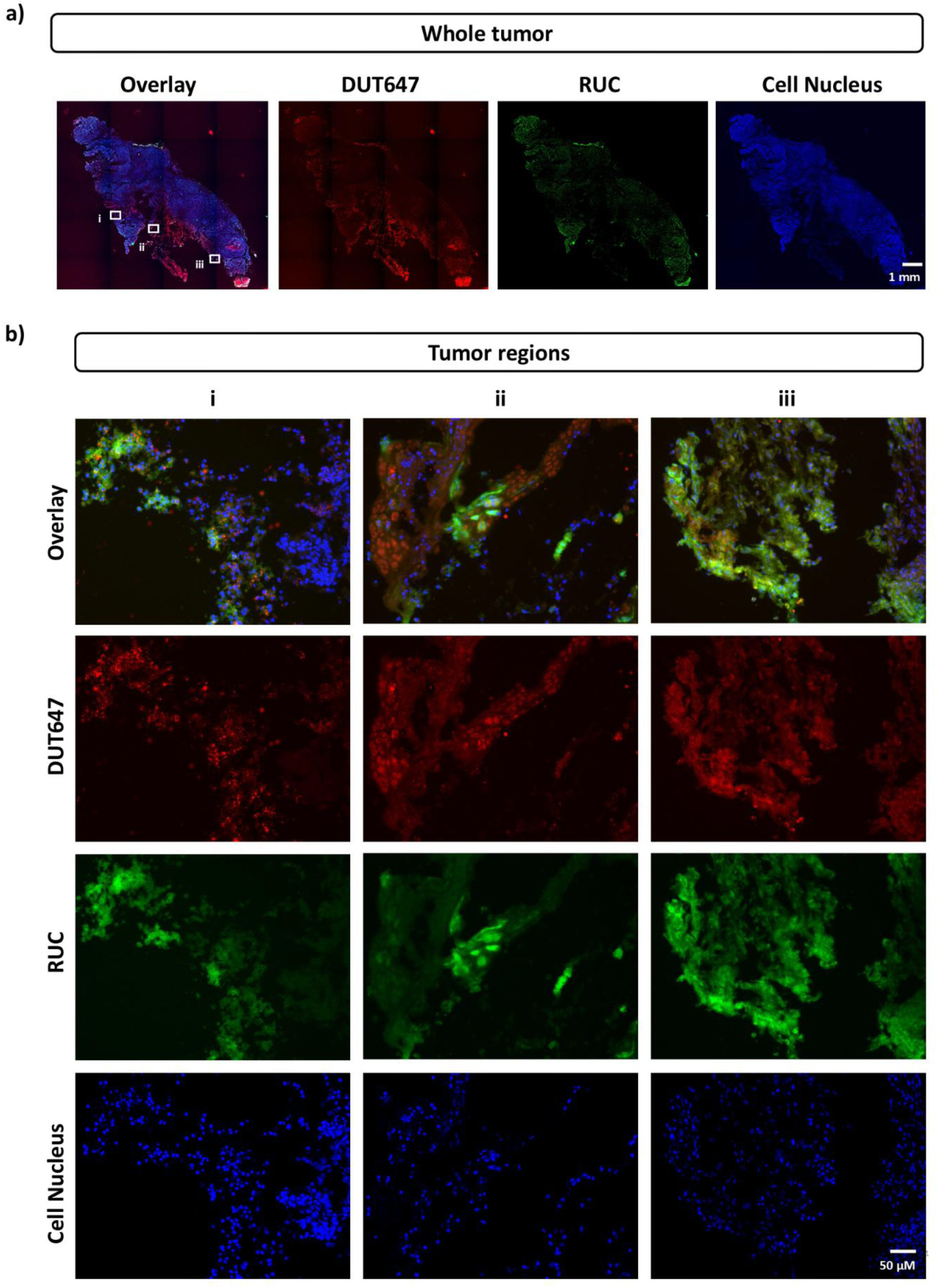
In vivo tumor uptake of RUC@[ZrO][FdUMP/DUT647] in a subcutaneous HT29 xenograft model 24 h post-injection. One athymic nude mouse was injected intravenously with 714 µg RUC@[ZrO][FdUMP/DUT647] in 0.9% NaCl. Microscopy images show the shell dye DUT647 of RUC@[ZrO][FdUMP/DUT647] (red), the core drug RUC of RUC@[ZrO][FdUMP/DUT647] (green) and cell nuclei (blue). **a)** Overview image of the tumor demonstrating heterogeneous uptake of RUC@[ZrO][FdUMP/DUT647] with high-uptake regions at the tumor rim. **b)** Three high-uptake regions from a), indicated as i, ii, and iii, showing different staining patterns of DUT647 and RUC and indicating efficient drug release from RUC@[ZrO][FdUMP/DUT647].

## 4. Discussion

### 4.1. High-drug loading RUC@[ZrO][FdUMP/DUT647] nanocarriers

We developed and evaluated RUC@[ZrO][FdUMP/DUT647] as a novel drug delivery system for combination chemotherapy in colorectal cancer. Using a solvent-antisolvent synthesis approach, we were able to encapsulate the lipophilic drug RUC together with the hydrophilic 5-FU metabolite FdUMP into a single nanocarrier with a drug load of 59 wt%. Considering that other nanocarriers, which are discussed below, typically show a drug load of 1 to 15 wt%, we consider this exceptionally high [46–48]. High drug-loading maximizes therapeutic efficacy while minimizing the amount of nanocarrier required for administration. Moreover, the simple three-step synthesis of RUC@[ZrO][FdUMP/DUT647] and the high drug load may simplify manufacturing compared with more complex multi-component formulations and could support future translational development. Complex formulations and low drug loading are among the major limiting factors that make clinical translation of nanocarriers challenging [47–49]. Consequently, only a limited number of studies have investigated nanocarriers co-delivering PARP inhibitors with other chemotherapeutics. For instance, Cavanagh et al. developed ROS-responsive polymeric nanocarriers encapsulating olaparib and doxorubicin, which demonstrated enhanced cytotoxicity in vitro compared to the free-drug combination [46]. However, drug loadings remained low at 3–3.7 wt% [46]. Nanoencapsulation further offers the advantage of delivering a predefined drug ratio to the tumor site. Therefore, Wang et al. first identified the most synergistic combination of niraparib and doxorubicin and subsequently synthesized a folate receptor-targeted liposome co-encapsulating both drugs at a synergistic ratio [50]. However, no biological evaluation of these liposomes was reported [50]. In addition, the efficient and stable encapsulation of lipophilic drugs, such as PARP inhibitors, which can freely diffuse through lipid bilayers, has proven challenging. Liang et al. addressed this limitation and reported the self-assembly of hydrophilic cisplatin with the PARP inhibitor olaparib together with heparin, demonstrating synergistic therapeutic efficacy in vivo and the ability to overcome chemoresistance in ovarian cancer models [51]. Still, 90 % of olaparib was released within 3 h in vitro, whereas we observed only 17.1% RUC release from RUC@[ZrO][FdUMP/DUT647] after 3 h in complete cell culture medium [51]. The unique core@shell structure of RUC@[ZrO][FdUMP/DUT647] enabled a drug load of 59 wt% and thereby addresses key translational limitations previously reported for PARP inhibitor–based multidrug nanocarrier systems. Furthermore, we demonstrated good nanocarrier stability of the lipophilic PARP inhibitor RUC.

### 4.2. Rapid cellular uptake of RUC@[ZrO][FdUMP/DUT647] nanocarriers

Using fluorescent imaging, we were able to monitor the subcellular fate of RUC@[ZrO][FdUMP/DUT647] after uptake. Microscopically, we observed an increasing co-localization of the shell-dye DUT647 and the intrinsically fluorescent core drug RUC after RUC@[ZrO][FdUMP/DUT647] treatment in vesicular structures. Since nanocarriers are typically taken up by endocytosis, we suggest that intact nanocarriers accumulate in endolysosomal compartments over time [52]. Specifically, ITC/TocP@ZrO(FdUMP] nanocarriers, which had the same core@shell structure, similar surface properties, and diameter as RUC@[ZrO][FdUMP/DUT647] were taken up by macropinocytosis [53]. In addition, we observed a clear RUC signal in the nucleus, indicating efficient release of the core drug RUC from the nanocarrier and subsequent binding of RUC to its target localization. Importantly, the stability of RUC@[ZrO][FdUMP/DUT647] in complete cell culture medium showed 47.3% free RUC after 24 h, and therefore, we expected some RUC to have passively crossed the cell membrane before nanocarrier uptake. Nevertheless, drug sequestration via nanocarrier accumulation in vesicular structures resulted in sustained nuclear levels of RUC.

The characterization of uptake kinetics via flow cytometry revealed that RUC@[ZrO][FdUMP/DUT647] uptake followed a similar kinetic to the free-drug cocktail of RUC and FdUMP, which was characterized by a steep increase in the first hour and reaching a plateau after 2 h. Other studies with [^18^F]- or [^14^C]-labeled RUC reported a similar increase in intracellular RUC within the first hour [54, 55]. Since the initial uptake of nanocarriers by macropinocytosis is already completed within the first minutes, it was not surprising to observe a similar uptake kinetic profile for RUC@[ZrO][FdUMP/DUT647] and the free-drug cocktail [56]. Both, RUC@[ZrO][FdUMP/DUT647] and the free RUC/FdUMP incubation, showed a further increase of the MFI between 6 and 24 h, which was absent when cells were incubated with RUC alone. We hypothesize that the presence of FdUMP leads to disruption of the cell membrane integrity [57], which enabled higher uptake of RUC.

In addition, we found that RUC@[ZrO][FdUMP/DUT647] treatment resulted in higher retention levels of RUC in HT29 cells, but not in HCT116 cells, when compared with the free-drug cocktail. This difference could be attributed to the fact that HT29 cells overexpress ATP-binding cassette (ABC) transporters and thereby, quickly externalize free drugs like RUC [58]. If RUC was encapsulated as RUC@[ZrO][FdUMP/DUT647], however, higher levels of RUC were retained intracellularly over time. Therefore, RUC@[ZrO][FdUMP/DUT647] nanocarriers may help increase intracellular retention of RUC in ABC transporter-high HT29 cells. This observation motivates future studies in dedicated drug-resistant models to evaluate whether the platform can mitigate resistance mechanisms. Nanoparticle-based drug delivery has already been proposed as a potential strategy to overcome ABC transporter-mediated drug resistance in colorectal cancer [59].

### 4.3. Synergistic cytotoxicity of nanocarrier-mediated co-delivery of RUC and FdUMP

RUC@[ZrO][FdUMP/DUT647] treatment resulted in similar cytotoxicity to the free-drug cocktail, and we identified the combination of RUC and FdUMP to be synergistic in colorectal cancer cells. In line with our findings, Falzacappa et al. identified synergism between RUC and 5-FU (pro-drug of FdUMP) in leukemic cells, while data from Augustine et al. suggested a greater-than-additive effect in HCT116 colorectal cancer cells [16, 17]. Our data showed that in HT29 cells, however, this combination effect was concentration-dependent because low and high doses exhibited synergism, while doses around the EC_50_ value had an additive or antagonistic effect. We hypothesize that at concentrations around the EC_50_ value, the FdUMP concentration was sufficiently high to induce S-phase arrest, which restricted RUC activity in the S- and subsequent G2-phase due to the greater cytotoxicity of FdUMP at these concentrations. At high doses, however, both agents exhibited a strong cytotoxic effect. In line with this, our cell cycle analysis confirmed that FdUMP induced greater cell cycle changes than RUC at equimolar concentrations. We found a significant increase in S-phase arrest after free RUC and FdUMP or RUC@[ZrO][FdUMP/DUT647] treatment, and fewer cells reaching the G2-phase. While FdUMP inhibits DNA synthesis and, thereby, causes S-phase arrest, RUC, like all PARP inhibitors, causes replication fork collapse and G2/M checkpoint activation, which is expected to induce S- and G2/M-phase arrest [15, 60]. Moreover, we found that RUC@[ZrO][FdUMP/DUT647] treatment was at least equally effective in inducing DNA DSB as the free-drug cocktail. RUC induced a greater increase in DNA DSB than FdUMP despite the lower cytotoxicity of RUC. RUC, like all PARP inhibitors, exerts its cytotoxicity by inducing DNA DSB, making it more effective at DNA DSB formation than FdUMP. Thus, RUC@[ZrO][FdUMP/DUT647] or the free-drug cocktail treatment holds potential as a radiosensitizing treatment, even in homologous recombination proficient HT29 cells. Overall, we conclude that RUC@[ZrO][FdUMP/DUT647] treatment resulted in S-phase arrest, which can largely be attributed to FdUMP from the shell, and an increase in DNA DSB, which was predominantly induced by the core drug RUC. These two molecular mechanisms act synergistically, as DNA DSB formation has been reported to be most pronounced following S-phase arrest after combination treatment of olaparib and 5-FU [16].

### 4.4. Efficient tumor uptake of RUC@[ZrO][FdUMP/DUT647] in HT29 xenograft

We further conducted a preliminary in vivo study, where one HT29-tumor-bearing mouse was injected with RUC@[ZrO][FdUMP/DUT647]. This proof-of-principle study showed efficient uptake of RUC@[ZrO][FdUMP/DUT647] in an HT29 subcutaneous tumor, as visualized by ex vivo fluorescence microscopy. Nanocarrier uptake was highly heterogeneous throughout the tumor, with the highest accumulation primarily at the tumor rim. This limited distribution is to be expected in subcutaneous tumor models, where vascularization is often restricted to the tumor surface and nanocarrier uptake is dependent on the enhanced permeability and retention (EPR) effect, whereby leaky tumor vasculature and impaired lymphatic drainage enable nanocarrier extravasation and retention [61, 62]. Nevertheless, the overlapping, but distinct staining patterns of the shell-dye DUT647 and the core drug RUC visualized efficient drug release from the nanocarrier. However, the in vivo uptake was only investigated in one animal and further validation is required to confirm tumor uptake and treatment efficacy. Despite our promising in vitro results, drug delivery of RUC@[ZrO][FdUMP/DUT647] remains to be investigated in vivo. Pharmacokinetics and biodistribution of RUC@[ZrO][FdUMP/DUT647] could be determined using fluorescence in vivo imaging or nuclear imaging after nanocarrier radiolabeling. Since RUC has a short emission wavelength, limiting its ability to penetrate tissue, and the concentration of DUT647 is very low per injection, multiple injections might be required for longitudinal fluorescence imaging in vivo.

## 5. Conclusions

In conclusion, we realized a novel high-drug loading nanocarrier design for combination chemotherapy by integrating the lipophilic PARP inhibitor rucaparib (RUC) with the hydrophilic 5-FU metabolite FdUMP in a single core@shell platform. RUC@[ZrO][FdUMP/DUT647] nanocarriers exhibited fast cellular uptake and triggered molecular responses comparable to the free-drug combination, including S-phase arrest and increased γH2AX foci, consistent with replication stress and DNA double-strand break formation. Drug–drug interaction analysis confirmed synergistic activity of the RUC/FdUMP pair in colorectal cancer cells. In drug efflux transporter-high HT29 cells, nanocarrier delivery increased intracellular retention of RUC relative to free drug, motivating future studies in dedicated drug-resistant models. Finally, our pilot in vivo experiment showed tumor-associated fluorescence signals 24 h post-injection, supporting further systematic evaluation of pharmacokinetics, biodistribution, efficacy, and safety in vivo.

## 6. Ethics approval

All animal experiments were conducted in accordance with the German Animal Welfare Act (Deutsches Tierschutzgesetz) under license ROB-55.2–2532.Vet_02-19–62 and ROB-55.2–2532.Vet_02-23–224 from the state government of upper Bavaria (Regierung von Oberbayern).

## 7. AI disclosure

During the preparation of this work the authors used ChatGPT, Grammarly, Gemini and Microsoft 365 Copilot TUM campus license in order to assist with language editing and proofreading. After using these tools, the authors reviewed and edited the content as needed and take full responsibility for the content of the published article.

## 8. Acknowledgments

We thank Deutsche Krebshilfe for funding within the collaborative CANACO project (Nr. 70114704). We thank the Preclinical Imaging Core Facility (PICTUM) at the Central Institute for Translational Cancer Research (TranslaTUM) and Markus Mittelhäuser and Natalie Röder for technical assistance.

## 9. Author declaration

Anna Erika Weber: Methodology (equal), investigation (equal), data curation (equal), formal analysis (equal), visualization (equal), writing original draft (equal), writing review and editing (equal)

Christian Ritschel: Methodology (equal), investigation (equal), data curation (equal), formal analysis (equal), visualization (equal), writing original draft (equal), writing review and editing (equal)

Susanne Kossatz: Conceptualization (equal), formal analysis (equal), writing review and editing (equal), supervision (equal), funding acquisition (equal)

Claus Feldmann: Conceptualization (equal), formal analysis (equal), writing review and editing (equal), supervision (equal), funding acquisition (equal)

