## Supplementary Information for "High Drug-Loading Rucaparib-FdUMP Nanocarriers for Colorectal Cancer Combination Therapy"

**<u>Content</u>**

**1. Analytical Techniques for Material Characterization**
**2. Synthesis of Nanocarriers**
**3. Material Characterization of Nanocarriers**
**4. In-vitro/in-vivo Studies Methodology**
**5. In-vitro/in-vivo Studies Supplementary Data**

### 1. Analytical Techniques for Material Characterization

**Dynamic light scattering (DLS)** was performed at room temperature in polystyrene cuvettes applying a Nanosizer ZS (Malvern Instruments, United Kingdom).

**Zeta potential measurements** were conducted using an automatic titrator MPT-2 attached to a Nanosizer ZS (Malvern Instruments, United Kingdom). Titrations were performed by addition of 0.1 M HCl or 0.1 M NaOH.

**Fourier-transformed infrared (FT-IR) spectroscopy** was performed on a Bruker Vertex 70 FT-IR spectrometer (Bruker, Germany). All samples were pestled and diluted with KBr (1 mg of sample in 300 mg of KBr) and pressed to pellets for transmission measurements.

**Elemental analysis (C/H/N/S analysis)** was performed via thermal combustion with an Elementar Vario Microcube device (Elementar, Germany) at a temperature of 1100 °C.

**Scanning electron microscopy (SEM)** was conducted with a Zeiss Supra 40 VP (Zeiss, Germany), equipped with a field-emission gun and a resolution of 1.3 nm (at 15 kV). Samples were prepared by placing small droplets of diluted aqueous suspensions on a silica wafer.

**Transmission electron microscopy (TEM)** and high-angle annular dark-field scanning transmission electron microscopy (HAADF-STEM) were conducted with a FEI Osiris microscope at 200 kV (FEI, The Netherlands). TEM samples were prepared by evaporating aqueous suspensions on amorphous carbon (Lacey-)films suspended on copper grids.

**Energy-dispersive X-ray spectroscopy (EDXS)** was measured at 200 kV electron energy with a FEI Osiris microscope (FEI, The Netherlands) that was equipped with a Bruker Quantax system (XFlash detector, Bruker, Germany). Quantification was performed with the FEI software package “TEM imaging and analysis” (TIA).

**X-ray powder diffraction (XRD)** was conducted on a Stadi-P diffractometer (Stoe, Germany) with Ge-monochromatized Cu-$K_{\alpha}$ radiation. Dried powder samples were fixed between Scotch tape and acetate paper.

**Thermogravimetry (TG)** was performed with a Netzsch STA 449 F3 (Netzsch, Germany) applying α-$Al_2O_3$ as crucible material and reference sample. The samples were heated under air to 1200 °C with a heating rate of 5 K/min.

**Optical spectroscopy (UV-Vis spectroscopy)** was performed with an UV2700 from Shimadzu (Japan). Nanocarrier suspensions were measured in UV-transparent polystyrene cuvettes in an integrating sphere in diffuse transmission geometry against the corresponding pure solvent as a reference.

**Photoluminescence (PL)** was recorded with a Horiba Jobin Yvon Spex Fluorolog 3 (Horiba Jobin Yvon, France) equipped with a 450 W Xe-lamp. Nanocarrier suspensions were measured in UV-transparent polystyrene cuvettes.

#### 2. Synthesis of Nanocarriers

**Starting materials.**

Rucaparib (RUC, MCE, USA, 99.8%), tocopherolphosphate disodium salt ($Na_2$(TocP), Sigma Aldrich, Germany, 97%), $ZrOCl_2 \times 8H_2O$ (Sigma Aldrich, Germany, 98%), DMSO (VWR, Germany, 99 %), disodium uridine monophosphate ($Na_2$(UMP), Sigma Aldrich, Germany, 98 %), trisodium citrate (Carl-Roth, Germany, 99%), DY$^{TM}$-647P1-aadUTP (DUT647, Dyomics, Germany), and disodium 5-fluoro-2′-deoxyuridine 5′-monophosphate ($Na_2$(FdUMP), Indagoo, Spain, 97%) were used as purchased.

**Synthesis**

*TocP-stabilized RUC nanocarriers*. 0.8 mg $Na_2$(TocP) (1.4 µmol, 0.2 eq.) were dissolved in 7 mL of ice water. Separately, 2.5 mg RUC (7.7 µmol, 1 eq.) were dissolved in 100 µL of DMSO and injected into the aforementioned solution. 3.5 mg $ZrOCl_2 \times 8H_2O$ (10.9 µmol, 1.4 eq.) were dissolved in 0.5 mL water and injected under ultrasonic treatment (Sonopuls HD 2070, Bandelin, Germany, 20 kHz, 70 W; 10 s, amplitude 100 %). Thereafter the suspension was adjusted to pH 7.0 by addition of 0.01 M NaOH. The TocP-stabilized RUC nanocarriers were centrifuged and redispersed- in/from water (pH 7.0) to remove excess starting materials and salts.

*RUC@[ZrO][UMP] core@shell nanocarriers.* The aforementioned TocP-stabilized RUC nanocarriers were intensely stirred and also sonicated (Bandelin, Germany, Sonopuls HD 2070,

20 kHz, 70 W; 10 s, amplitude 100%) at pH 7.0, while a solution of 3.5 mg $Na_2$(UMP) (9.5 µmol, 1.2 eq.) in 0.5 mL of water was injected. The suspension was again centrifuged and redispersed in water (pH 7.0). Thereafter, a solution of 2.8 mg $ZrOCl_2 \times 8H_2O$ (8.7 µmol, 1.1 eq.) in 2 mL of water and 3.3 mg $Na_2$UMP (9.0 µmol, 1.2 eq.) dissolved in 2 mL of water were simultaneously added via syringe pump (2 mL/h, HLL LA120, Germany). Finally, the suspension of the RUC@[ZrO][UMP] core@shell nanocarriers was centrifuged/redispersed twice from/in water to remove excess starting materials and salts. For long-term storage (2-3 month), the nanocarriers (2.7 mg/mL) were redispersed in a solution of trisodium citrate (5 mg/mL).

*RUC@[ZrO][FdUMP] core@shell nanocarriers.* These nanocarriers were prepared similar to the aforementioned RUC@[ZrO][UMP] core@shell nanocarriers. Instead of $Na_2$(UMP), however, $Na_2$(FdUMP) was applied, using 3.1 mg (8.4 µmol, 1.1 eq.) for the injection step and 3.0 mg (8.1 µmol, 1.1 eq.) for slow addition via syringe pump. The molar amount of $Na_2$(FdUMP) was slightly reduced in comparison to $Na_2$(UMP). Mainly driven by the high cost of $Na_2$(FdUMP), the molar excess in relation to $ZrOCl_2 \times 8H_2O$ was reduced, which still results in colloidally stable suspensions.

*DUT647-labeled RUC@[ZrO][UMP] or RUC@[ZrO][FdUMP] core@shell nanocarriers.* For fluorescence labeling, a small amount of DUT647 was added to the [ZrO][UMP] or [ZrO][FdUMP] shell of the nanocarriers. To this concern, 40 nmol of DUT647 were dissolved in 100 µL of water and added to the $Na_2$(UMP) or $Na_2$(FdUMP) solution in the above synthesis recipe.

### 3. Material Characterization of Nanocarriers

*TocP-stabilized RUC nanocarriers (RUC NC)*

The as-formed TocP-stabilized RUC nanocarriers (RUC NC) are colloidally stable on a time scale of some hours (*see main paper: Figure 1*). A longer period of stirring at room temperature, however, resulted in an increased turbidity of the suspensions, which originated from an increase of the particle size from about 20 nm (Figure S1a-c) to about 90 nm (Figure S1d-f) due to Ostwald ripening. The TocP-stabilized RUC nanocarriers can be stabilized further when increasing the amount of TocP. Here, the TocP concentration was kept as low as possible, and the RUC nanocarriers were stabilized by a [ZrO][UMP] or [ZrO][FdUMP] shell.

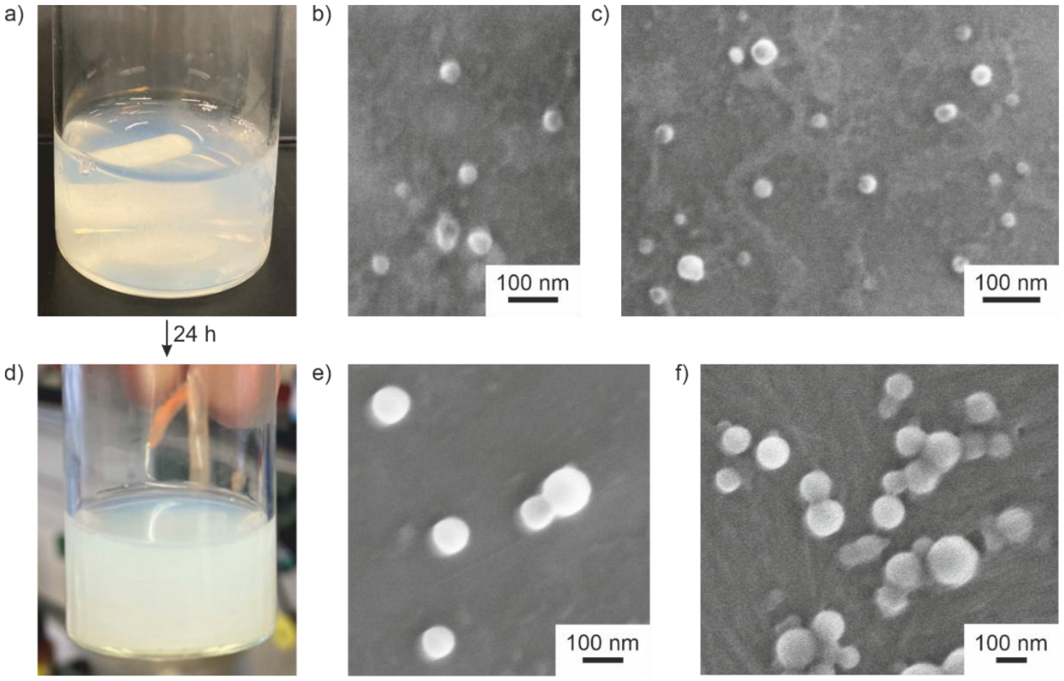


**Figure S1. Particle growth of TocP-stabilized RUC nanocarriers over longer periods of time. a)** Photo of as-prepared suspension. **b+c)** SEM images with particle diameters of about 20 nm. **d)** Photo of suspension after 24 hours of stirring at room temperature. **e+f)** SEM images with particle diameters of about 90 nm.

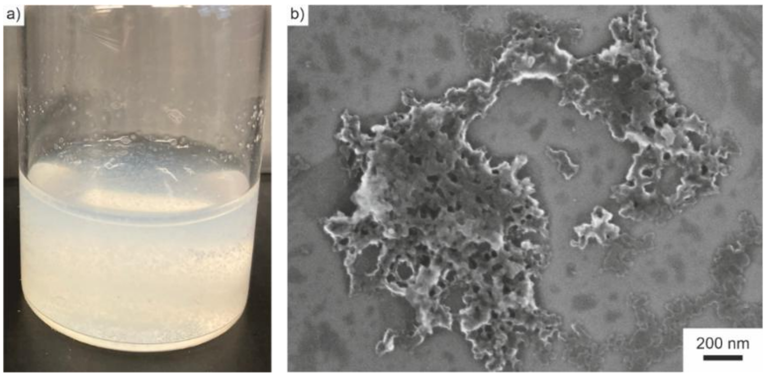


**Figure S2. Colloidal stability of TocP-stabilized RUC nanocarriers after addition of $ZrOCl_2 \times 8H_2O$. a)** Photo of suspension. **b)** SEM image of aggregating nanocarriers.

After the addition of $ZrOCl_2 \times 8H_2O$, the TocP-stabilized RUC nanocarriers are colloidally less stable due to lower negative surface charging with the $[ZrO]^{2+}$ cations binding to the phosphate groups of TocP on the nanocarrier surface (*see main paper: Figure 1*). Therefore, SEM images show significant agglomeration of the nanocarriers (Figure S2), which is addressed by simultaneous addition of $Na_2$(UMP) or $Na_2$(FdUMP) in the course of the synthesis.

Fourier-transform infrared (FT-IR) spectra of TocP-stabilized RUC nanocarriers exhibit vibrations that prove the presence of RUC and TocP (Figure S3a). The most characteristic vibrations relate to $\nu$(O–H) at 3430 cm$^{-1}$ originating from RUC (3431 cm$^{-1}$ due to keto-enol tautomerism) as well as $\nu$(N–H) and $\nu$(C–H) at 3298, 3183, 3055, 2926, and 2868 cm$^{-1}$. These vibrations are also observed in the reference spectra of RUC (3273, 3171, 3036, 2940, 2805 cm$^{-1}$) and TocP (2928, 2897, 2870, 2845 cm$^{-1}$). In addition, $\nu$(C=O) of RUC (1616 cm$^{-1}$) occurs in the spectra of the nanocarriers (1611 cm$^{-1}$). Finally, the broad absorption at 3690-3080 cm$^{-1}$ is attributed to $\nu$(O–H) of water adsorbed on the nanocarrier surface.

X-ray powder diffraction (XRD) indicate the TocP-stabilized RUC nanocarriers to be amorphous (Figure S3b). This finding is in difference to pure RUC as a starting material, which is highly crystalline. The non-crystallinity of RUC in the nanocarriers can be attributed to the small size of the nanocarriers as well as to the presence of the non-polar surfactant tails of TocP that are located between the RUC molecules.

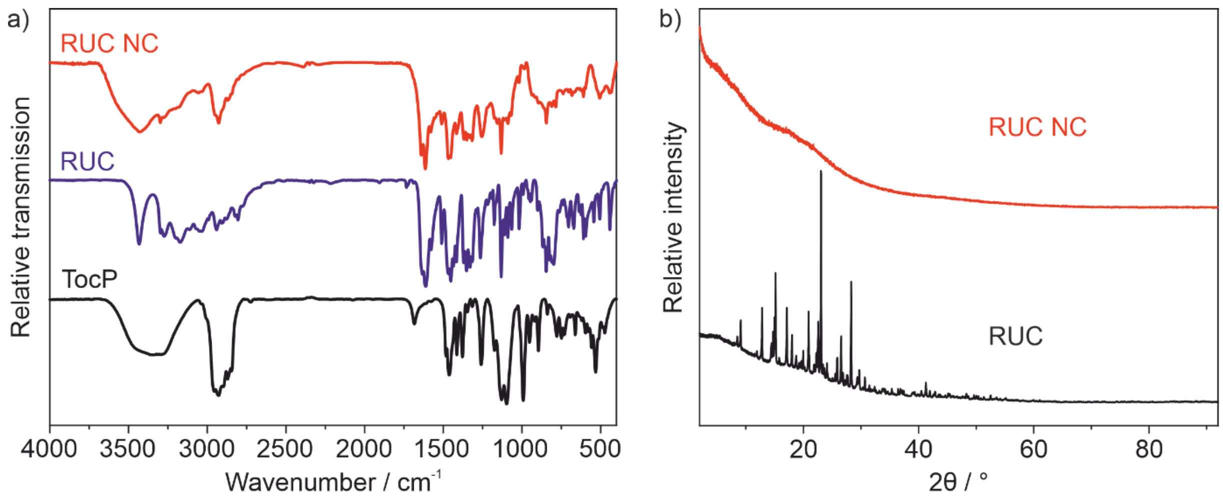


**Figure S3. Composition of TocP-stabilized RUC nanocarriers. a)** FT-IR spectrum (with RUC and $Na_2$(TocP) as references). **b)** XRD (RUC as a reference).

The composition of the TocP-stabilized RUC nanocarriers was analyzed by total organic combustion with thermogravimetry (TG) and elemental analysis (EA). TG shows a two-step decomposition (Figure S4a) with a first decomposition at 40-260 °C and 13.4 % mass loss, which can be attributed to an evaporation of surface-adsorbed water and DMSO. A second decomposition at 260-750 °C with 44.2% can be related to the decomposition of all organic constituents.

After accounting for the surface-adsorbed solvents (7.3 wt% water and 6.1 wt% DMSO), the C/H/N/S contents determined by elemental analysis (EA) (i.e., 4.6 wt% N, 35.0 wt% C,

4.7 wt% H, 2.5 wt% S) were corrected to 5.3 wt% N, 39.0 wt% C, 4.7 wt% H, and 0.0 wt% S (Table S1).

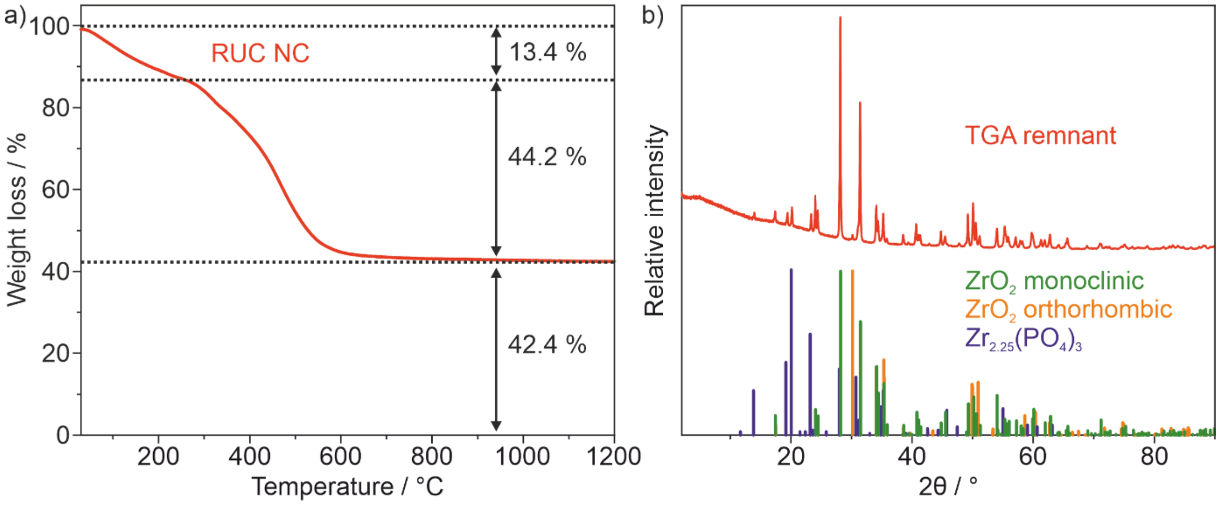


**Figure S4. Thermal analysis of TocP-stabilized RUC nanocarriers. a)** TG (in air). **b)** XRD of the TG residue (references: monoclinic $ZrO_2$/ICSD-No. 94886, orthorhombic $ZrO_2$/ICSD-No. 67004, $Zr_{2.25}(PO_4)_3$/ICDD-No. 00-064-0782).

**Table S1. C/H/N/S/Zr contents of TocP-stabilized RUC nanocarriers (dried in vacuum, $10^{-3}$ mbar, 20 °C, 1 h) according to EA and TG.**

| | **C wt-% (EA)** | **H wt-% (EA)** | **N wt-% (EA)** | **S wt-% (EA)** | **Zr wt-% (TG)** |
|---|---|---|---|---|---|
| Experimental data | 35.0 | 4.7 | 4.6 | 2.5 | <31.4 |
| Corrected for surface adhered DSMO (6.1 wt%) | 35.3 | 4.5 | 4.9 | 0.0 | <33.4 |
| Corrected for surface adhered DSMO (6.1 wt%) and water (7.3 wt%) | 38.1 | 4.0 | 5.3 | 0.0 | <36.2 |
| Calculated data | 39.0 | 4.3 | 5.3 | 0.0 | 29.3 |

Based on the nitrogen content, the RUC content of TocP-stabilized RUC nanocarriers can be deduced to 40.8 wt-%, as RUC is the sole nitrogen-containing component in the nanocarriers. After subtracting the RUC-associated carbon content, the remaining carbon content can be attributed to TocP, resulting in a TocP content of 13.6 wt%. Considering the 2.9 wt% of $[ZrO]^{2+}$ cations, which are bound to the $[TocP]^{2-}$ anions, the remaining 42.7 wt% can be attributed to $ZrO(OH)_2$. The presence of $ZrO(OH)_2$ is likely due to the excess use of $ZrOCl_2$ during synthesis and the subsequent neutralization of the reaction mixture with NaOH solution. Reducing the amount of $ZrOCl_2$ leads to colloidally non-stable RUC@[ZrO][UMP] or RUC@[ZrO][FdUMP] nanocarrier suspensions at the end of the synthesis (Figure S5). It is highly probable that an excess of $[ZrO]^{2+}$ is essential for achieving a positive surface charge, facilitating the ionic binding of the required UMP/FdUMP to the nanocarrier surface and ensuring the formation of colloidally stable suspensions.

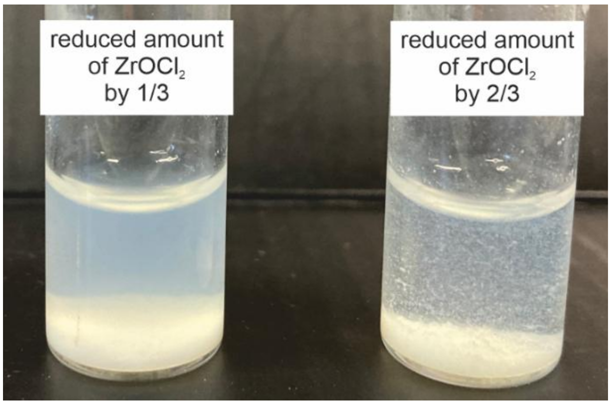


**Figure S5. Photos of colloidally unstable suspensions of RUC@[ZrO][UMP] nanocarriers synthesized with a reduced amount of $ZrOCl_2 \times 8H_2O$.**

XRD of the thermal residue of the TG analysis after heating to 1200 °C reveals intense Bragg reflections corresponding to monoclinic $ZrO_2$ and weaker reflections of orthorhombic $ZrO_2$ (Figure S4b). Additionally, $Zr_{2.25}(PO_4)_3$ is detected. This confirms the presence of $[ZrO]^{2+}$ in the TocP-stabilized RUC nanocarriers. While the Zr content in the nanocarriers cannot be precisely determined from the TGA and XRD data due to the presence of different Zr-containing phases, $ZrO_2$ is identified as a significant thermal residue in the TGA, providing an estimate for the Zr content. Assuming only $ZrO_2$ is present in the thermal residue, the Zr content in the nanocarriers is 31.3 wt% (solvent-corrected 36.2 wt%) (Table S1). Since $Zr_{2.25}(PO_4)_3$, which is also present, has a lower Zr weight fraction than $ZrO_2$, the actual Zr content is likely lower than 36.2 wt%.

In summary, the TocP-stabilized RUC nanocarriers are described by the formula $(RUC)_{13}@([ZrO][TocP])_3((ZrO(OH)_2)_{30})$ with 40.9 wt-% RUC, which is consistent with 40.8 wt-% obtained by C/H/N/S analysis. The calculated Zr content is 29.3 wt%, which also agrees with the experimental Zr content of 31.3 wt-% (solvent-corrected 36.2 wt-%).

In addition to elemental analysis and total-organic content determined based on powder samples, the supernatant of suspensions after separation of the TocP-stabilized RUC nanocarriers via centrifugation was analyzed by UV-Vis spectroscopy. This approach provides additional information about the nanocarrier composition. As reference measurements can only be performed with solutions (i.e. RUC in DMSO, TocP in water), the transmittance of the supernatant was analyzed for quantitative evaluation, rather than analyzing the nanocarriers themselves directly. This method ensures greater accuracy. Comparing the spectra of intact nanocarrier suspensions with those of reference solutions can be error-prone, primarily due to the stronger scattering of light by the nanocarrier suspensions.

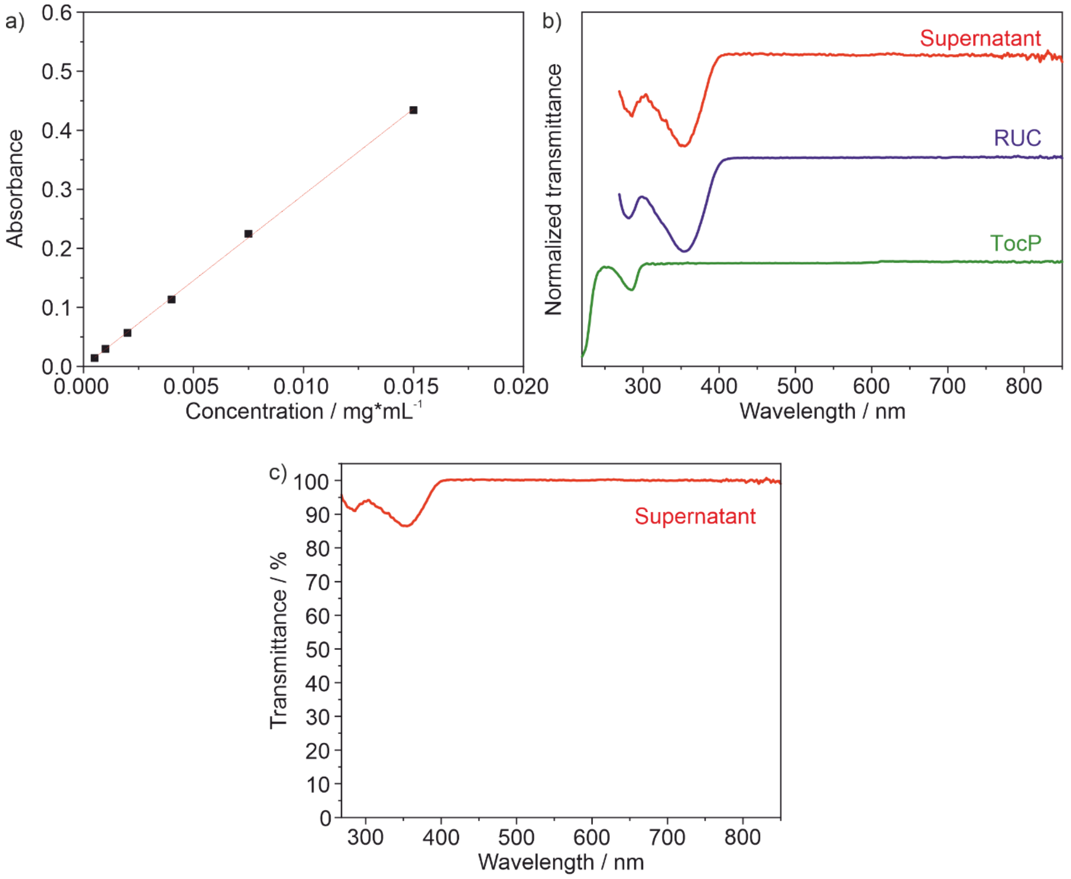


**Figure S6. Photometric analysis of the supernatant after separation of TocP-stabilized RUC nanocarriers by centrifugation. a)** Photometric calibration curve of RUC at $\lambda_{max}$ = 355 nm (solutions in DMSO). **b)** Normalized transmission spectra of the supernatant (dried and dissolved in DMSO) and RUC (in DMSO) as well as of TocP (in water) as references. **c)** Absolute transmission spectrum of the supernatant (dried and dissolved in DMSO).

To determine the RUC content in TocP-stabilized RUC nanocarriers, a calibration curve was determined based on UV-Vis spectra with RUC solutions (in DMSO) at different concentrations, using the absorption maximum of RUC ($\lambda_{max}$ = 355 nm) (Figure S6a). The supernatant obtained after centrifugation following the addition of $ZrOCl_2$ is water-based, making a direct comparison with the calibration curve measured in DMSO prone to significant error. Therefore, the supernatant was dried overnight (approximately 18 h at 60 °C), and the resulting solid residue was dissolved in DMSO. The transmittance spectrum of this DMSO-based solution is shown in Figure S6b and matches the reference spectrum of RUC, with no overlaps observed with the TocP spectrum. This allows for a quantitative determination of the RUC content in the supernatant.

The absorbance at $\lambda_{max}$ = 355 nm for the DMSO-based supernatant solution was determined (Figure S6c). To ensure the absorbance fell within the linear range of the RUC calibration curve, the DMSO-based supernatant solution was diluted. Specifically, the dried supernatant was dissolved in 10 mL of DMSO, and 50 µL of this solution was further diluted with 1950 µL of DMSO. Using this method, the total amount of RUC in the supernatant was calculated to be 0.9 mg. Since 2.5 mg of RUC was used for the synthesis, it was determined that 1.6 mg of RUC (4.9 mmol) remained in the nanocarriers. The yield of dried TocP-stabilized RUC nanocarriers per synthesis was approximately 4 mg, resulting in a RUC content of 40.0 wt% in the nanocarriers, according to UV-VIS spectroscopy. This value is in good agreement with the composition calculation based on TGA and EA (RUC content 40.8 wt%).

*RUC@[ZrO][UMP] and RUC@[ZrO][FdUMP] core@shell nanocarriers*

To modify the TocP-stabilized RUC nanocarriers with a [ZrO][UMP] or [ZrO][FdUMP] shell, first of all, the surface of the TocP-stabilized RUC nanocarriers is functionalized by UMP/FdUMP via injection of a solution of $Na_2$(UMP)/$Na_2$(FdUMP) (*see Section 2: Synthesis of Nanocarriers*). This intermediate is analyzed in the following.

TG of RUC@[ZrO][UMP] nanocarriers with a thin [ZrO][UMP] shell shows a two-step decomposition with a first step (60-205 °C) due to the evaporation of adsorbed water (8.5 wt-%) and residual DMSO (0.7 wt%) and a second step (205-650 °C, 56.3 wt-%) due to total organics combustion (Figure S7a). The thermal residue (34.5 wt%) was identified by XRD as mixture of $ZrO_2$ and $Zr_{2.25}(PO_4)_3$ (Figure S7b). Assuming the residue consists solely of $ZrO_2$, the theoretical maximum zirconium content is 25.5 wt% (or 28.1 wt% when corrected for solvent content). However, due to the presence of $Zr_{2.25}(PO_4)_3$ in the residue, the actual zirconium content is expected to be slightly lower.

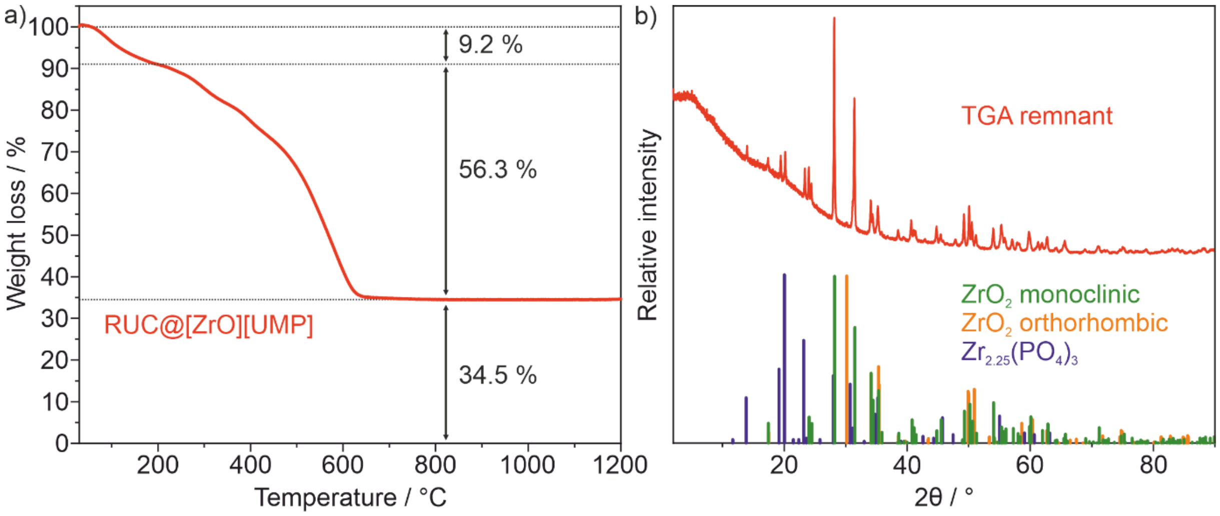


**Figure S7. Thermal analysis of RUC@[ZrO][UMP] core@shell nanocarriers with a thin [ZrO][UMP] shell. a)** TG (in air). **b)** XRD of thermal TG residue (monoclinic $ZrO_2$/ICSD-No. 94886, orthorhombic $ZrO_2$/ICSD-No. 67004, $Zr_{2.25}(PO_4)_3$/ICDD-No. 00-064-0782 as references).

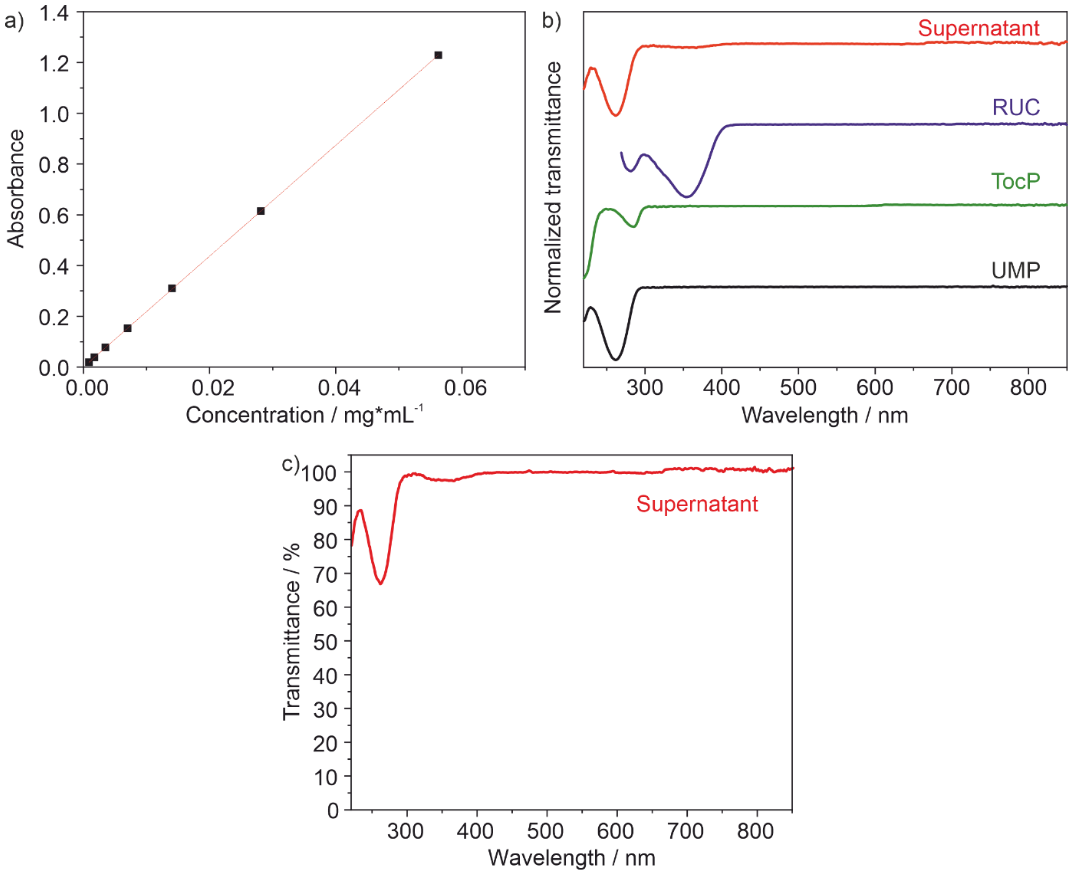


**Figure S8. UV-VIS analysis of the reaction supernatant after separation of RUC@[ZrO][(UMP)] nanocarriers by centrifugation. a)** Photometric calibration curve of UMP (solution in water). **b)** Normalized transmittance spectra of the supernatant (water-based)

with RUC (dissolved in DMSO), TocP (dissolved in water), and UMP (dissolved in water) as references. **c)** Absolute transmittance spectrum of the diluted supernatant.

Quantifying the amount of UMP-bound in the nanocarrier shell is challenging, as UMP has a nitrogen content (8.6 wt%) comparable to that of RUC (7.7 wt%) and does not introduce any new elements not already present in the initial nanocarriers. However, since UMP binds exclusively to $[ZrO]^{2+}$ cations already incorporated within the nanocarrier structure, an increase in the overall nitrogen content is expected following its addition. This expected increase in nitrogen content in the RUC@[ZrO][UMP] nanocarriers, relative to the TocP-stabilized RUC nanocarriers, was confirmed by elemental analysis (EA).

EA of the RUC@[ZrO][UMP] nanocarriers yielded the following elemental compositions: 6.5 wt% N, 37.3 wt % C, 4.9 wt% H, and 0.3 wt% S. After correcting for the presence of adsorbed water (8.5 wt%) and residual DMSO (0.7 wt%), the values adjust to 7.1 wt% N, 40.8 wt% C, 4.3 wt % H, and 0.0 wt% S (Table S2). In comparison, the TocP-stabilized RUC nanocarriers exhibited nitrogen contents of 4.6 wt%, or 5.3 wt% after solvent correction. Thus, the incorporation of UMP results in an approximate increase of 2.0 wt% in nitrogen content, supporting its successful integration into the nanocarrier shell.

**Table S2. C/H/N/S/Zr contents of RUC@[ZrO][UMP] nanocarriers with a thin [ZrO][UMP] shell (dried in vacuum, $10^{-3}$ mbar, 20 °C, 1 h) according to EA and TG.**

| | **C wt-% (EA)** | **H wt-% (EA)** | **N wt-% (EA)** | **S wt-% (EA)** | **Zr wt-% (TG)** |
|---|---|---|---|---|---|
| Experimental data | 37.3 | 4.9 | 6.5 | 0.3 | <25.5 |
| Corrected for surface adhered DSMO (0.7 wt-%) | 37.3 | 4.9 | 6.5 | 0.0 | <25.7 |
| Corrected for surface adhered DSMO (0.7 wt-%) and water (8.5 wt-%) | 40.8 | 4.3 | 7.1 | 0.0 | <28.1 |
| Calculated data | 39.2 | 4.1 | 6.1 | 0.0 | 24.0 |

Furthermore, the thermal residue remaining after heating the RUC@[ZrO][UMP] nanocarriers to 1200 °C was 34.5 wt% (38.0 wt% after solvent correction), which is notably lower than that of the TocP-stabilized RUC nanocarriers (42.4 wt%; corrected to 49.0 wt%). This decrease indicates a higher overall organic content in the UMP-containing nanocarriers. The experimental data align well with a proposed composition of

$(RUC)_{13}$@($[ZrO][TocP])_3$($[ZrO][UMP]_7$)($ZrO(OH)_2)_{22}$, corresponding to an RUC-to-UMP ratio of 13:7 (Table S2).

To further quantify the amount of UMP bound to the nanocarrier surface, UV-Vis spectroscopy was performed on the reaction supernatant obtained by centrifugation. A calibration curve for UMP was established at its absorption maximum $\lambda_{max} = 261$ nm (*see Figures S13a*). Since the absorption properties of the nanocarriers may differ from those of pure reference compounds, as previously discussed for RUC@[ZrO][TocP] nanocarriers, analyzing the reaction supernatant provides a more reliable basis for quantitative evaluation in this case as well. Moreover, the transmittance spectrum of the TocP solution partially overlaps with that of UMP, making direct calculations from a RUC@[ZrO][TocP] nanocarrier suspension prone to inaccuracies (Figures S8b). The transmittance spectrum of the supernatant closely resembles that of pure UMP, showing a prominent absorption band at $\lambda_{max} = 261$ nm and only weak absorption at $\lambda = 355$ nm, which can be attributed to RUC. No significant absorption associated with TocP is observed. These findings confirm that a quantitative determination of UMP in the supernatant is feasible (Figures S8b).

To quantify the UMP content in the supernatant, the transmittance at $\lambda_{max} = 261$ nm was measured for a diluted sample (50 µL of supernatant diluted with 1950 µL of water). Using the absorbance value and the known dilution factor, the amount of unbound UMP in the supernatant was calculated. The analysis indicates that 2.5 mg of UMP remain in solution, while 1.0 mg (corresponding to 2.7 mmol) are incorporated into the nanocarriers to form the shell of the RUC@[ZrO][UMP] nanocarriers (Figures S8c). This result aligns well with the observed increase in nanocarrier yield during synthesis, where approximately 5 mg of RUC@[ZrO][UMP] powder was obtained, an increase of ~1 mg compared to the yield from RUC@[ZrO][TocP] nanocarriers.

To verify that FdUMP behaves similarly to UMP, UV-Vis measurements were performed on the RUC@[ZrO][FdUMP] nanocarriers, where FdUMP replaced UMP during synthesis. A calibration curve for FdUMP was established at $\lambda_{max} = 269$ nm using transmittance measurements at various concentrations (Figures S9a). The transmittance spectrum of the reaction supernatant closely matches that of FdUMP, with the addition of a low-intensity absorption at $\lambda = 355$ nm, which can be attributed to RUC (Figures S9b). This allows for a quantitative determination of the amount of FdUMP in the supernatant. By measuring the diluted supernatant, the total amount of FdUMP remaining in solution was found to be 2.1 mg (Figures S9c). Given that 3.1 mg of FdUMP was used in the synthesis, this implies that 1.0 mg

(2.7 mmol) of FdUMP remains bound to the nanocarriers. These results confirm that FdUMP behaves similarly to UMP in this synthesis process.

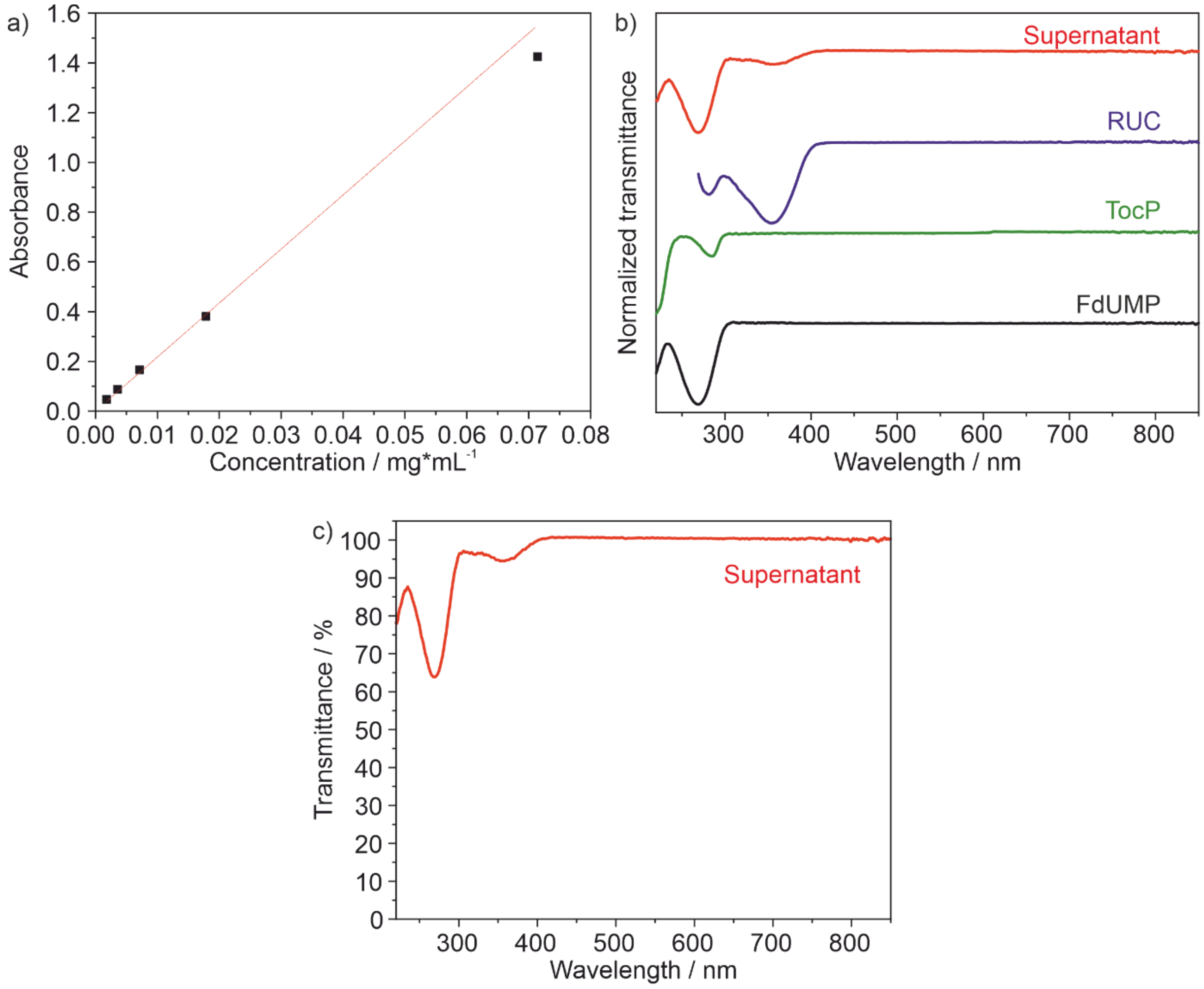


**Figure S9. UV-VIS analysis of the supernatant after separation of RUC@[ZrO][FdUMP] nanocarriers by centrifugation. a)** Photometric calibration curve of FdUMP (solution in water). **b)** Normalized transmittance spectra of the supernatant (water-based) and RUC (dissolved in DMSO) as well as TocP (dissolved in water) as references. **c)** Transmittance spectrum of the diluted supernatant.

Subsequent to the surface functionalization of the TocP-stabilized RUC nanocarriers with a first thin [ZrO][UMP] shell (*see Section 2: Synthesis of Nanocarriers*), the thickness of the [ZrO][UMP] shell was increased by simultaneous slow addition of $Na_2$(UMP) and $ZrOCl_2$ (*see Section 2: Synthesis of Nanocarriers*). These final RUC@[ZrO][UMP] core@shell nanocarriers are characterized in the following.

Powder samples of RUC@[ZrO][UMP] core@shell nanocarriers exhibit a yellow color, which originates from RUC and, thus, already indicates its presence (Figure S10a). FT-IR spectra show the vibrations that can be attributed to RUC, TocP, and UMP (Figure S10b). Specifically, P=O vibrations (988 $cm^{-1}$) originate from TocP (991 $cm^{-1}$) and UMP (980 $cm^{-1}$). N–H (3268, 3105 $cm^{-1}$) and C–H vibrations (2955, 2926, 2868 $cm^{-1}$) can be assigned to RUC (3273, 3171, 3036, 2940, 2805 $cm^{-1}$), UMP (2924, 2876 $cm^{-1}$), and TocP (2928, 2870 $cm^{-1}$). Furthermore, C=O vibrations from RUC (1611 $cm^{-1}$) and UMP (1678 $cm^{-1}$) are present in the spectrum (1690, 1616 $cm^{-1}$). A broad absorption at 3700-3000 $cm^{-1}$ can be associated with O–H vibrations and the presence of water. Water remains adsorbed on the surface of the nanocarriers due to the water-based synthesis. The broad absorption partially overlaps with the O–H vibrations of the hydroxyl groups of RUC (3431 $cm^{-1}$ due to keto-enol tautomerism) and UMP (3534 $cm^{-1}$). X-ray powder diffraction (XRD) indicates the as-prepared RUC@[ZrO][UMP] core@shell nanocarriers to be amorphous (Figure S10c).

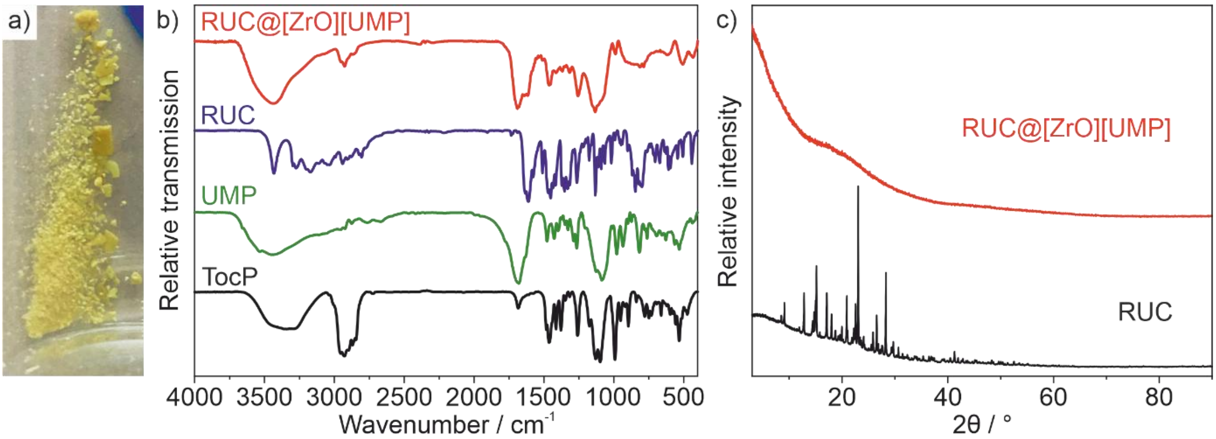


**Figure S10. Composition of RUC@[ZrO][UMP] core@shell nanocarriers. a)** Photo of powder sample. **b)** FT-IR spectrum (with RUC, $Na_2$(UMP), $Na_2$(TocP) as references). **c)** XRD (with RUC as a reference).

To further characterize the composition, particularly the amounts of RUC and UMP in the RUC@[ZrO][UMP] nanocarriers (or RUC and FdUMP in the RUC@[ZrO][FdUMP] nanocarriers), the analytical methods EA, TGA, and UV-VIS spectroscopy were used. The analysis was based on characterization data obtained for surface functionalization of the TocP-stabilized RUC nanocarriers with a first thin [ZrO][UMP] shell. It is assumed that the composition of the core nanocarriers remains unchanged during the subsequent addition of $ZrOCl_2$ and UMP solutions. Therefore, the observed changes in composition can be explained solely by the additional binding of $[ZrO]^{2+}$ cations and $[UMP]^{2-}$ anions to the surface.

TGA analysis was performed on dried powder samples ($10^{-3}$ mbar, 1 h) of RUC@[ZrO][UMP] nanocarriers and shows a three-stage decomposition. In the first stage (50-

200 °C), DMSO and water molecules (10.5 wt%) that were adsorbed on the surface of the nanocarriers evaporate. The combustion of the organic components of the nanocarriers results in a second weight loss (200–600 °C) of 43.2%. A third decomposition stage (600–920 °C) can be attributed to the formation of pyrophosphate through a condensation reaction of phosphate anions, leading to a weight loss of 1.8% (Figure S11a). Overall, the decomposition results in a weight loss of 55.5% at 1200 °C. The thermal residue was analyzed by powder XRD and consists of $ZrO_2$, $Zr_{2.25}(PO_4)_3$, and $ZrP_2O_7$ (Figure S11b).

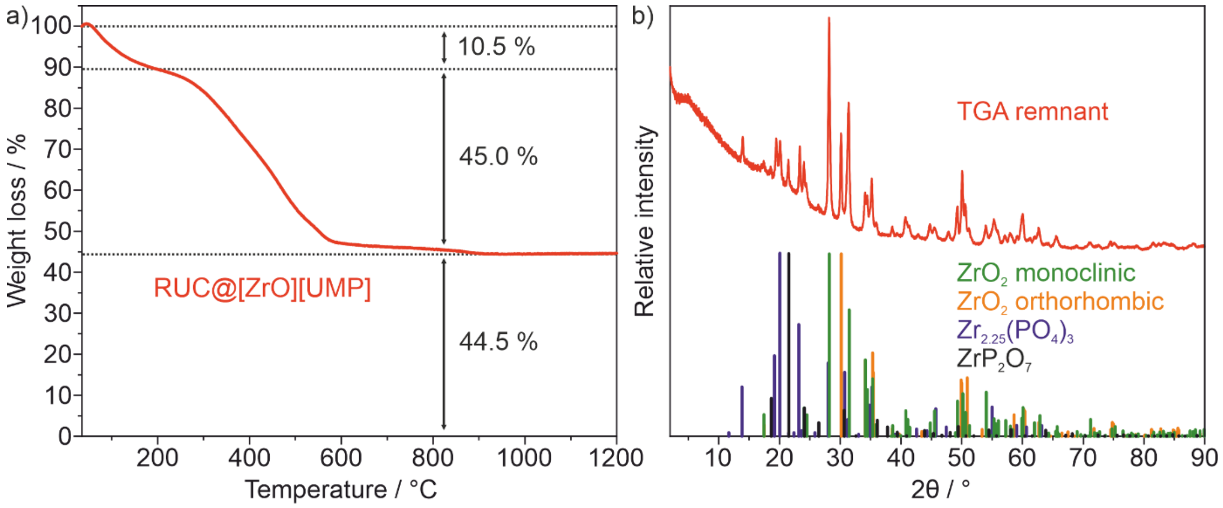


**Figure S11. Chemical composition of RUC@[ZrO][UMP] nanocarriers. a)** TGA under air (30–1200 °C). **b)** Powder XRD of the thermal TGA residue with monoclinic $ZrO_2$ (ICSD No. 94886), orthorhombic $ZrO_2$ (ICSD No. 67004), and $Zr_{2.25}(PO_4)_3$ (ICDD No. 00-064-0782) as references.

Furthermore, the weight percentages of the elements C, N, H, and S were determined using EA. The measured values (5.4 wt% N, 29.1 wt% C, 3.9 wt% H, and 0.2 wt% S) were corrected for the amount of adsorbed water (10.0 wt%) and DMSO (0.5 wt%), resulting in values of 6.0 wt% N, 32.3 wt% C, 3.1 wt% H, and 0.0 wt% S. Because of the presence of various Zr-based combustion products identified in the diffraction pattern, the exact Zr content could not be determined from the data. However, a maximum value of 32.9 wt% (or solvent-corrected 36.8 wt%) can be estimated assuming the thermal residue consists solely of $ZrO_2$. Since this assumption does not reflect the actual composition (Figure S11b), and $Zr_{2.25}(PO_4)_3$ and $ZrP_2O_7$ contain less Zr than $ZrO_2$, the actual value must be lower (Table S3). Based on these data, determining the exact drug content of the synthesized nanocarriers remains challenging. Nonetheless, a proposed composition of the form

$RUC_{13}$@([ZrO][TocP])$_3$([ZrO][UMP])$_7$(ZrO(OH)$_2$)$_{22}$@([ZrO][UMP])$_9$ is in good agreement with the measured data (Table S3).

**Table S3. C/H/N/S/Zr contents of final RUC@[ZrO][UMP] nanocarriers (dried in vacuum, $10^{-3}$ mbar, 20 °C, 1 h) according to EA and TG.**

| | C wt-% (EA) | H wt-% (EA) | N wt-% (EA) | S wt-% (EA) | Zr wt-% (TG) |
|---|---|---|---|---|---|
| Experimental data | 29.1 | 3.9 | 5.4 | 0.2 | <32.9 |
| Corrected for surface adhered DSMO (0.5 wt-%) | 29.1 | 3.9 | 5.4 | 0.0 | <33.1 |
| Corrected for surface adhered DMSO (0.5 wt-%) and water (10.0 wt-%) | 32.3 | 3.1 | 6.0 | 0.0 | <36.8 |
| Calculated data | 35.8 | 3.8 | 6.2 | 0.0 | 23.3 |

UV-Vis spectroscopy measurements were conducted to quantify the amount of UMP bound to the surface of the nanocarriers and to verify the EA results. The UV-VIS spectrum of the supernatant obtained through centrifugation during the synthesis shows a strong absorption at $\lambda_{max}$ = 261 nm, which can be attributed to the presence of UMP (Figure S12a). Additionally, a weak absorption at $\lambda$ = 355 nm was observed, attributed to RUC. RUC is slightly soluble in water and can be released in small amounts from the nanocarriers; furthermore, a few nanocarriers may remain in the supernatant after centrifugation. Overall, there was no significant spectral overlap with TocP or RUC at $\lambda$ = 261 nm, allowing for a quantitative analysis of UMP in the supernatant. 50 µL of the supernatant, from a total of 11 mL, were diluted with 1950 µL of water, and a UV-VIS spectrum was recorded (Figure S12b). Using a calibration curve of UMP (*see Figure S8*) and the transmittance of the diluted supernatant at $\lambda_{max}$ = 261 nm (76.86%), the total amount of UMP in the supernatant was calculated to be 2.2 mg. Since 3.3 mg of UMP were used for the synthesis, 1.1 mg (3.0 mmol) remain in the nanocarriers. Overall, based on the UV-VIS analysis of the supernatant after each reaction step, 2.1 mg of UMP (5.7 mmol) and 1.6 mg of RUC (4.9 mmol) are integrated into the RUC@[ZrO][UMP] nanocarriers. This results in a molar ratio of UMP to RUC of 1.2:1, which is in good agreement with the ratio in the proposed composition based on TGA and EA ($RUC_{13}$@([ZrO][TocP])$_3$([ZrO][UMP])$_7$(ZrO(OH)$_2$)$_{22}$@([ZrO][UMP])$_9$ with a UMP-to-RUC ratio of 1.2:1).

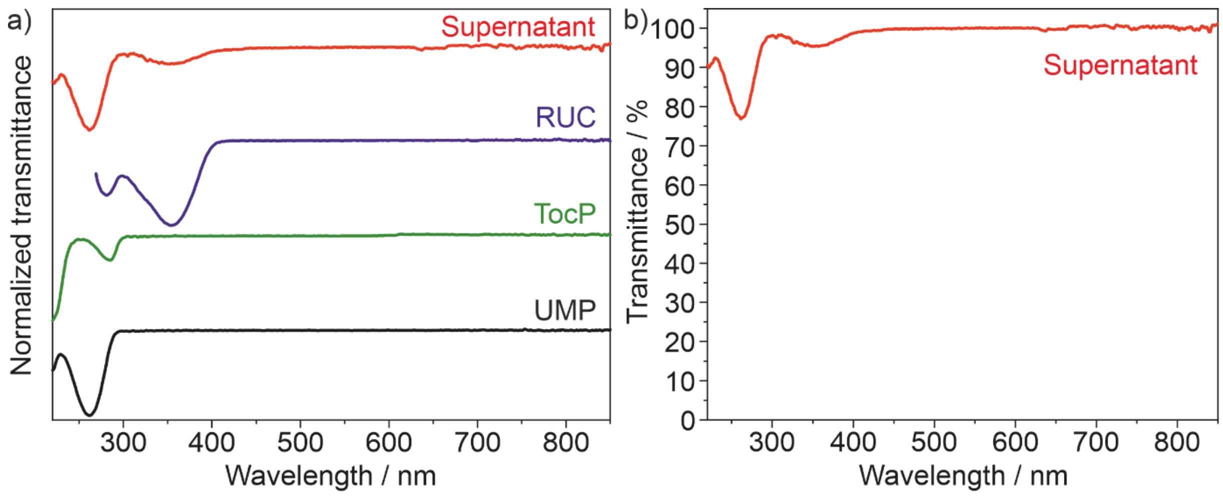


**Figure S12. UV-VIS analysis of the supernatant after centrifugation to separate the RUC@[ZrO][UMP] nanocarriers. a)** Normalized transmittance spectra of the supernatant (water-based) as well as RUC (dissolved in DMSO), TocP (dissolved in water) and UMP (dissolved in water) as references. **b)** Absolute transmittance spectrum of the diluted supernatant.

The described UV-VIS analysis of the supernatant was also conducted for the FdUMP-based synthesis. In this case, no significant overlap was observed between the spectrum of the supernatant at $\lambda = 269$ nm and those of TocP or RUC (Figure S13a). Therefore, a quantitative analysis of FdUMP in the supernatant can be performed. The measured transmittance of 80.75% for the diluted supernatant corresponds to a total of 1.9 mg of FdUMP in the supernatant (Figure S13b). Consequently, 1.1 mg (3.0 mmol) of the 3.0 mg of added FdUMP is incorporated into the shell of the nanocarriers.

The FdUMP-to-RUC ratio can now be calculated as well from the UV-VIS analysis. After completing all synthesis steps, the nanocarriers contain 1.6 mg of RUC (4.9 mmol) and 2.1 mg of FdUMP (5.7 mmol), corresponding to an FdUMP-to-RUC ratio of 1.2:1. This ratio is comparable to the UMP-to-RUC ratio in the RUC@[ZrO][UMP] nanocarriers, demonstrating the similarity between FdUMP and UMP in this synthesis process.

All characterization methods collectively suggest a nanocarrier composition that can be described by the formula $RUC_{13}$@([ZrO][TocP])$_3$([ZrO][UMP])$_7$(ZrO(OH)$_2$)$_{22}$@([ZrO][UMP])$_9$ respectively $RUC_{13}$@([ZrO][TocP])$_3$([ZrO][FdUMP])$_7$(ZrO(OH)$_2$)$_{22}$@([ZrO][FdUMP])$_9$. The synthesized nanocarriers show a drug load of 26.2 wt% RUC and 32.3 wt% FdUMP, thus, 58.5 wt% drug load in total relative to the mass of the nanocarriers.

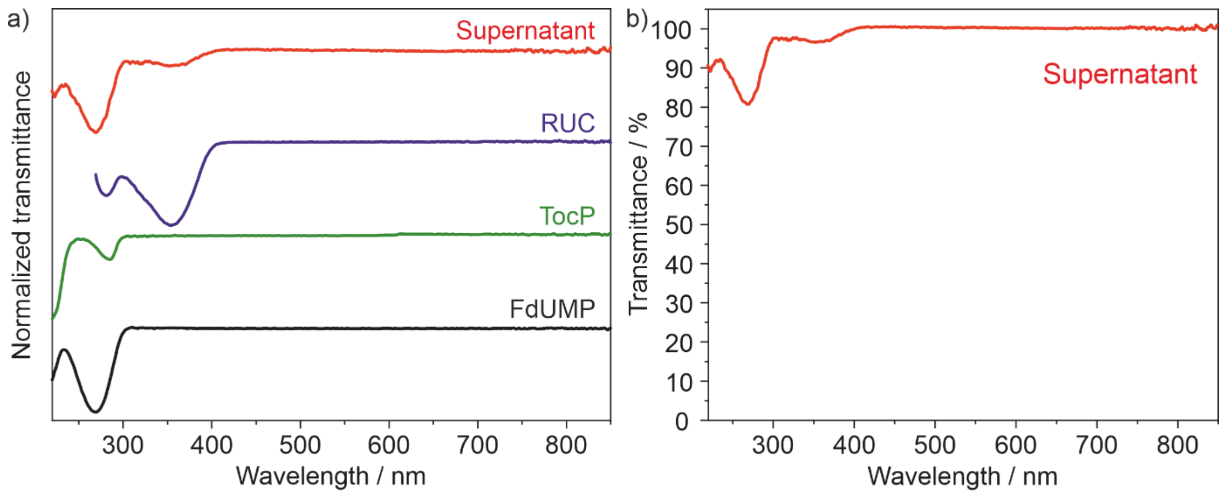


**Figure S13. UV-VIS analysis of the supernatant after centrifugation to separate the RUC@[ZrO][FdUMP] nanocarriers. a)** Normalized transmittance spectra of the supernatant (water-based) as well as RUC (dissolved in DMSO), TocP (dissolved in water) and FdUMP (dissolved in water) as references. **b)** Absolute transmittance spectrum of the diluted supernatant.

To track the synthesized nanocarriers with optical imaging both in vitro and in vivo, fluorescent dye labeling is useful. Since RUC exhibits intrinsic blue fluorescence, no additional labeling was needed for the core of the RUC@[ZrO][UMP] nanocarriers.[14] DY 647P1-dUTP (DUT647), a fluorescent dye functionalized with a uridine triphosphate group, was used to label the shell. During synthesis, DUT647 can be incorporated into the nanocarriers alongside UMP or FdUMP anions. Due to its high luminescence intensity, only small amounts of DUT647 (40 nmol) were required for labeling (Figure S14). This resulted in the formation of bluish nanocarriers (the blue color is caused by DUT647), which, when excited by UV light ($\lambda_{Ex,max}$ = 360 nm), exhibited both blue emission (RUC-based, $\lambda_{Em,max}$ = 483 nm) and red emission (DUT647-based, $\lambda_{Em,max}$ = 681 nm) (Figure S14). The red emission can also be excited by orange-red light ($\lambda_{Ex,max}$ = 653 nm).

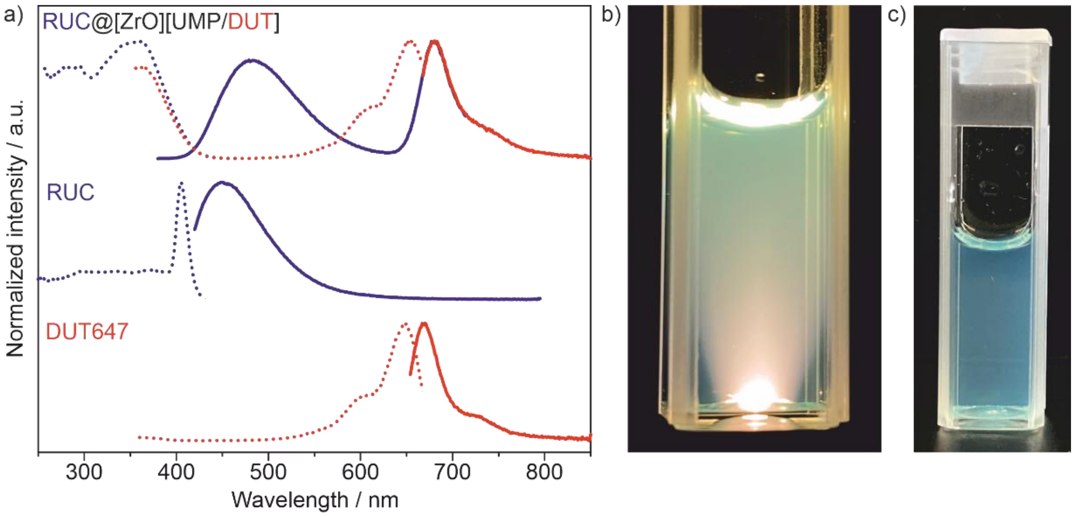


**Figure S14. Fluorescence labeling of RUC@[ZrO][UMP] nanocarriers with DUT647. a)** Excitation and emission spectra, including reference spectra of RUC and DUT647. **b)** Photo of a suspension under excitation with a cold light lamp. **c)** Photo of a suspension in daylight.

UV-VIS absorbance spectra of the RUC@[ZrO][UMP/DUT647] and the RUC@[ZrO][FdUMP/DUT647] nanocarriers showed absorption bands for RUC (λ = 355 nm), UMP (λ = 261 nm), and FdUMP (λ = 269 nm) (Figure S15), confirming the presence of the respective drug components. An additional absorption band (λ = 660 nm) relates to DUT647, used as a fluorescent marker in the nanocarrier shell, as previously described.

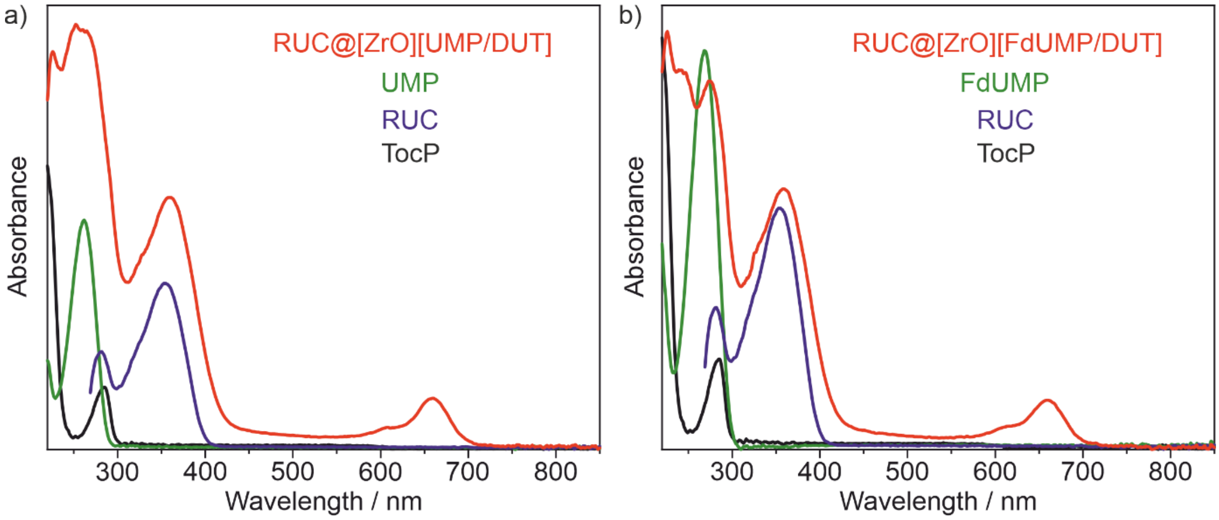


**Figure S15. UV-VIS absorbance spectra of a) RUC@[ZrO][UMP/DUT647] and b) RUC@[ZrO][FdUMP/DUT647] nanocarriers compared with the reference spectra of UMP, RUC, and TocP.** The nanocarrier shell is labeled with DUT647 resulting in an additional absorption band at 660 nm.

To evaluate the release of RUC from the RUC@[ZrO][UMP] core@shell nanocarriers, fixed aliquots of the as-prepared nanocarrier suspensions were poured in centrifuge tubes, and the RUC@[ZrO][UMP] core@shell nanocarriers were separated from the initial suspension medium (water) by centrifugation (Figure 3). The solid residue was then redispersed in the respective medium, in which the RUC release was studied. As a reference, the centrifuged RUC@[ZrO][UMP] core@shell nanocarriers were treated with DMSO for complete release of all RUC. The RUC content was then determined by UV-VIS spectroscopy and represents 100% of RUC load.

To evaluate the release of RUC from the nanocarriers, the centrifuged RUC@[ZrO][UMP] core@shell nanocarriers were redispersed and continuously stirred in various media at room temperature (20 °C) and or at 37 °C. After different time intervals (5 min, 1 h, 3 h, 6 h, 24 h), the remaining nanocarriers were again centrifuged and the solid residue extracted with DMSO. The change in concentration relative to the aforementioned reference was determined by UV-Vis spectroscopy (Figure 3).

### 4. In vitro/in vivo Studies Methodology

**Cell cultivation**

The human colon carcinoma cell lines HCT116 (ATCC, CCL-247) and HT29 (ATCC, HTB-38) were cultivated in McCoy's 5A (modified) medium (Gibco, #26600023) with 10% (v/v) heat inactivated FBS (Gibco, #A5256801) and 1% (v/v) P/S (Gibco) at 37°C in a humidified atmosphere of 95% and 5% (g/v) $CO_2$. All cell lines were regularly tested for mycoplasma and authenticated by PCR-single-locus-technology.

**Fluorescence microscopy of nanocarrier uptake**

To visualize cellular nanocarrier uptake, 10,000 HT29 cells were seeded per well on a poly-l-lysine-coated 8-well chamber slide (Ibidi, #80824). After overnight incubation, cells were treated with 5.65 µg/mL RUC@[ZrO][FdUMP/DUT647] nanocarriers (containing 5 µM RUC (RUC, MCE, USA, 99.8%) and 5.7 µM FdUMP (Indagoo, Spain, 97%) or 5 µM RUC and 5.7 µM FdUMP as a control for 0.5 h, 4 h or 24 h at 37°C. Cell nuclei were stained by adding Nuclear Green LCS1 (AAT Bioquest, #ABD-17540) to a final concentration of 8 µM for the last 20 min of treatment. Afterward, cells were washed three times with PBS and fixated with 4 % PFA in PBS for 8 min at room temperature. Finally, cells were washed with PBS and mounting medium (ROTI®Mount FluorCare, HP19.1, Roth) was added. Microscopy images were taken using an EVOS M7000 microscope (Thermo Fisher Scientific) and processed with

FIJI. The RUC signal was recorded in the DAPI cube ($\lambda_{ex}$ = 357/44 nm, $\lambda_{em}$ = 447/60 nm), the Nuclear Green LCS1 in the GFP cube ($\lambda_{ex}$ = 470/22 nm, $\lambda_{em}$ = 525/50 nm), and the nanocarrier shell dye DUT647 signal of RUC@[ZrO][FdUMP/DUT647] in the Cy5 cube ($\lambda_{ex}$ = 628/40 nm, $\lambda_{em}$ = 685/40 nm).

**Flow cytometric assessment of nanocarrier uptake**

To quantify cellular nanocarrier uptake, the RUC signal inside cells was measured with flow cytometry. Therefore, 100,000 HCT116 or HT29 cells were seeded per well on a 24-well plate (VWR, #734-2325). Cells were incubated for a total amount of 48 h and cells were treated for the last 5 min, 10 min, 30 min, 1 h, 2 h, 4 h, 6 h, and 24 h with 0.2 µg/mL RUC@[ZrO][FdUMP/DUT647] nanocarriers (containing 0.2 µM RUC and 0.2 µM FdUMP), 0.2 µM free RUC and 0.2 µM free FdUMP or 0.2 µM free RUC. Next, cells were trypsinized, transferred to 5 mL flow cytometry tubes and kept on ice. Cells were washed twice with flow buffer (2% fetal calf serum (w/v) (Gibco) in PBS) and resuspended in 15 nM of SYTOX Green dead cell stain (Invitrogen, #S7020). Experiments were performed on a NovoCyte flow cytometer (Agilent, 2060R) in three biological replicates. Cell clumps and debris were excluded through gating in the forward and side scatter. Gating was done with the software NovoExpress (Version 1.6.0). The SYTOX Green dead cell stain was detected in the FITC channel ($\lambda_{ex}$ = 488 nm, $\lambda_{em}$ = 530/30 nm), and gating was performed to exclude dead cells. The intracellular signal intensity of RUC was measured in the Pacific Blue channel ($\lambda_{ex}$ = 405 nm, $\lambda_{em}$ = 445/45 nm) and the DUT647 signal in the APC channel ($\lambda_{ex}$ = 637 nm, $\lambda_{em}$ = 675/30 nm).

**Flow cytometric assessment of intracellular RUC retention**

The intracellular retention of free or nanocarrier-delivered RUC was determined by measuring the RUC signal in cells at certain timepoints after treatment using flow cytometry. Therefore, 30,000 HCT116 or HT29 cells were seeded per well on a 24-well plate (VWR, #734-2325) and incubated overnight. The cells were treated for 4 h with 0.2 µg/mL RUC@[ZrO][FdUMP/DUT647] nanocarriers (containing 0.2 µM RUC and 0.2 µM FdUMP), 0.2 µM free RUC and 0.2 µM free FdUMP, or 0.2 µM RUC, followed by a 10 min wash in cell culture medium at room temperature and the addition of fresh medium for a post-incubation period of 10 min, 30 min, 1 h, 2 h, 4 h, 6 h, 24 h or 30 h. For analysis, cells were trypsinized and transferred to 5 mL flow cytometry tubes on ice. Cells were washed twice with flow buffer (2% fetal calf serum (w/v) (Gibco) in PBS) and resuspended in 15 nM of SYTOX Green dead cell stain (Invitrogen, #S7020). Experiments were performed on a NovoCyte flow cytometer

(Agilent, 2060R) in three biological replicates. Cell clumps and debris were excluded through gating in the forward and side scatter. Gating was done with the software NovoExpress (Version 1.6.0). The SYTOX Green dead cell stain was detected in the FITC channel ($\lambda_{ex}$ = 488 nm, $\lambda_{em}$ = 530/30 nm), and gating was performed to exclude dead cells. The intracellular signal intensity of RUC was measured in the Pacific Blue channel ($\lambda_{ex}$ = 405 nm, $\lambda_{em}$ = 445/45 nm), and the mean RUC signal per cell was normalized to the mean RUC signal after 4 h treatment for each biological repetition.

**Evaluation of free drug and nanocarrier cytotoxicity**

Cytotoxicity of RUC@[ZrO][FdUMP/DUT647] nanocarriers was assessed and compared to free drug treatment with RUC and/or FdUMP using the AlamarBlue HS (Invitrogen, #A50101) cell viability assay. Additionally, drug-free nanocarriers with the composition [ZrO][UMP/DUT647] were tested. In a 96-well plate (VWR, #732-3737), 1,500 HT29 or 2,000 HCT116 cells were seeded per well. After 24 h incubation, cells were treated for 72 h with varying treatment concentrations and afterward incubated with 10% AlamarBlue HS solution for 3 h. Subsequently, fluorescence was measured on a BioTek Synergy HT plate reader ($\lambda_{ex}$ = 540/35 nm, $\lambda_{em}$ = 590/20 nm), and the percentage of cell viability was calculated in relation to untreated cells. On the x-axis, the total concentration of all drugs in the treatment solution was plotted. For example, 30 µM referred to i) 30 µM RUC, ii) 30 µM FdUMP, iii) 10 µM RUC and 20 µM FdUMP (1:2), iv) 20 µM RUC and 10 µM FdUMP (2:1), or v) 15 µM RUC and 15 µM FdUMP (1:1). Each experiment was carried out three times with three technical replicates. The mean values of each biological repetition were plotted for each concentration in GraphPad Prism 10 using nonlinear regression analysis (Dose-response – Inhibition; [Inhibitor] vs. response (three parameters)). $EC_{50}$ (the half-maximal effective concentration) values described total drug concentrations in the treatment solution and were calculated from the plotted mean values of each biological repetition in GraphPad Prism 10 using nonlinear regression analysis (Dose-response – Inhibition; [Inhibitor] vs. response (three parameters)).

Based on the cell viability data, the combined effect of RUC and FdUMP (ratio 1:1) was determined to be synergistic, additive or antagonistic by the combinations index computed in the CompuSyn software (Version 1.0).

**Analysis of cell cycle arrest after treatment**

Distribution of the cell cycle in response to RUC@[ZrO][FdUMP/DUT647] or free drug treatment was assessed with flow cytometry. For this purpose, 60,000 HCT116 or HT29 cells were seeded per well on a 24-well plate (VWR, #734-2325) and incubated for 48 h. Thereafter, cells were incubated with either 1.13 μg/mL RUC@[ZrO][FdUMP/DUT647] (containing 1 μM RUC and 1.1 μM FdUMP), 1 μM RUC, 1 μM FdUMP, 1 μM RUC and 1 μM FdUMP or complete cell culture medium for 24 h. Cells were trypsinized, and transferred to 5 mL flow cytometry tubes on ice. Next, cells were washed with flow buffer (2% fetal calf serum (w/v) (Gibco) in phosphate-buffered saline (PBS) and fixated with 70% ethanol for 30 min at 4°C. Afterward, cells were washed twice, and 5 μg RNAse A (New England Biolabs) was added to 500 μL of cells in flow buffer. In a last step, one drop of propidium iodide Ready Flow Reagent (Invitrogen, R37169) was added, and measurements were performed on a NovoCyte flow cytometer (Agilent, 2060R). Gating and cell cycle analysis were performed with the software NovoExpress (Version 1.6.0). Cell clumps and debris were excluded using the forward and side scatter and propidium iodide stain was detected in the PE/Texas Red channel ($\lambda_{ex}$ = 488 nm, $\lambda_{em}$ = 615/20 nm). For cell cycle analysis, the percentage of cells in the G1-, S- and G2-phase was calculated using the Watson model and manually adjusted if necessary.

**Evaluation of DNA damage through the formation of γH2AX foci**

DNA damage was evaluated by fluorescent microscopy of γH2AX foci. Therefore, 10,000 HCT116 or HT29 cells were seeded per well on a poly-l-lysine-coated 8-well chamber slide (Ibidi, #80824). Cells were incubated overnight and treated with 5.65 μg/mL RUC@[ZrO][FdUMP/DUT647] (containing 5 μM RUC and 5.7 μM FdUMP), 5 μM RUC, 5 μM FdUMP, 5 μM RUC and 5 μM FdUMP or complete cell culture medium for 24 h. Thereafter, cells were washed three times in complete cell culture medium and fixated with 4% paraformaldehyde (Otto Fischar GmbH & Co. KG), followed by permeabilisation with 0.1% Triton X-100 (Roth). Blocking was performed using 3% bovine serum albumin (BSA) (Roth) for 1 h. Foci of γH2AX were stained by incubating cells with phospho-histone H2A.X Ser139 antibody (1:500, Thermo Fisher Scientific, #MA5-27753) in 3% BSA (Roth) and 0.1% Tween-20 (Roth) for 1 h at room temperature. Cells were washed three times in PBS, followed by incubation with goat anti-mouse IgG antibody Alexa Fluor 488 (1:2000, Thermo Fisher Scientific, #A11001) in 3% BSA (Roth) for 1 h at room temperature and three washing steps in PBS. Finally, cell nuclei were stained with 2 μL/mL Hoechst 33342 (Thermo Fisher Scientific). Microscopy was done with an EVOS M7000 microscope (Thermo Fisher Scientific), and images were processed in FIJI. Foci of γH2AX were recorded with the GFP cube ($\lambda_{ex}$ = 470/22 nm, $\lambda_{em}$ = 525/50 nm) and the nuclear Hoechst signal in the DAPI cube ($\lambda_{ex}$ = 357/44 nm, $\lambda_{em}$

= 447/60 nm). Quantification of γH2AX foci was performed with the software CellProfiler (Version 4.2.8) and the "Speckle counting" pipeline.

**Animal experiments**

One female seven-week-old athymic nude mouse (Crl:NU/NCr-Foxn1nu, Charles River Laboratories, Sulzfeld) was subcutaneously injected with $4 \times 10^6$ HT29 human colon carcinoma cells in 1:1 McCoy's 5A (modified) medium (Gibco) and Matrigel (Corning, #11543550). When the tumor reached ca. 300 $mm^3$, 714 µg RUC@[ZrO][FdUMP/DUT647] in 0.9% NaCl were intravenously injected into the mouse, and the animal was euthanized 24 h post-injection. The tumor was directly embedded in an optimal cutting temperature compound (Sakura), frozen, and cryosectioned into 10 µM sections. Sections were incubated in 4 µM Nuclear Green LCS1 (AAT Bioquest, #ABD-17540) for 30 min at room temperature and fixated with 4% paraformaldehyde (Otto Fischar GmbH & Co. KG). Microscopy images were taken with an EVOS M7000 microscope (Thermo Fisher Scientific) and processed with FIJI. The RUC signal of RUC@[ZrO][FdUMP/DUT647] was recorded with the DAPI cube ($\lambda_{ex}$ = 357/44 nm, $\lambda_{em}$ = 447/60 nm), the Nuclear Green LCS1 with the GFP cube ($\lambda_{ex}$ = 470/22 nm, $\lambda_{em}$ = 525/50 nm), and the shell dye DUT647 signal of RUC@[ZrO][FdUMP/DUT647] with the Cy5 cube ($\lambda_{ex}$ = 628/40 nm, $\lambda_{em}$ = 685/40 nm). As a control, the same staining and microscopic procedure were done on a subcutaneous C4-2 xenograft in a 25-week old male severe combined immunodeficient mice (SCID) mouse (Charles River Laboratories, Sulzfeld), that was used in another experiment without RUC@[ZrO][FdUMP/DUT647] injection.

**Statistical Analysis**

Graphs and statistical analysis were computed in GraphPad Prism (Version 10.0.3). All graphs presented mean values ± SD. Statistical significance was tested using the Mann-Whitney U test, with a p-value < 0.05 considered statistically significant.

### 5. In vitro/in vivo Supplementary Data

While measuring the intracellular RUC levels following RUC@[ZrO][FdUMP/DUT647], RUC, and RUC and FdUMP treatment (*see main paper: Figure 3*), the morphology of HCT116 and HT29 cells changed from 6 h to 24 h if FdUMP was present (Figure S15). Cells were treated with 0.2 µM free RUC or 0.2 µM free FdUMP and 0.2 µM free RUC. The forward and side scatter detectors showed that most cells were larger and more granular after 24 h treatment with RUC and FdUMP, while no changes were visible after 24 h RUC single treatment or 6 h treatment of RUC and FdUMP (Figure S15). Together with the increased intracellular RUC levels observed between 6 and 24 h of RUC and FdUMP or RUC@[ZrO][FdUMP/DUT647] treatment (*see main paper: Figure 3*), these data suggest that the cell membrane integrity is disrupted by FdUMP.

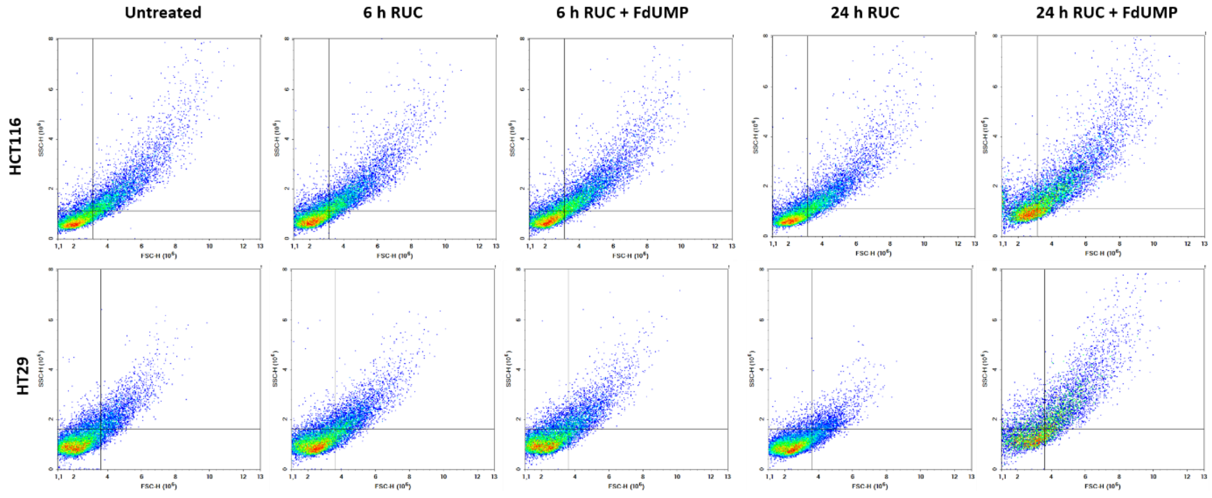


**Figure S15. Flow cytometric analysis of HCT116 and HT29 cell morphology after 6 h and 24 h of RUC or RUC and FdUMP treatment.** Cells were treated with 0.2 µM RUC or 0.2 µM RUC and 0.2 µM FdUMP.

To verify that the observed cytotoxic effects (*see main paper: Figure 4*) were not attributed to the solvent in the treatment solutions, HCT116 and HT29 viability was assessed with an AlamarBlue HS assay after incubation with the highest solvent concentrations of the tested treatments for 72 h (Figure S16). Compared to untreated cells, HCT116 and HT29 viability did not change after incubation with the RUC solvent (0.2 % DMSO for 100 µM RUC), the FdUMP solvent (5 % water for 127 µM FdUMP), the combination of RUC and FdUMP solvents (0.2 % DMSO and 4 % water for 100 µM RUC and 100 µM FdUMP), or the RUC@[ZrO][FdUMP/DUT647] solvent (773 µM trisodium citrate in 4.5 % water for Rucaparib@[ZrO][FdUMP] containing 100 µM RUC and 114 µM FdUMP) (Figure S16). Therefore, the observed cytotoxic effects (*see main paper: Figure 4*) were not influenced by the treatment solvents.

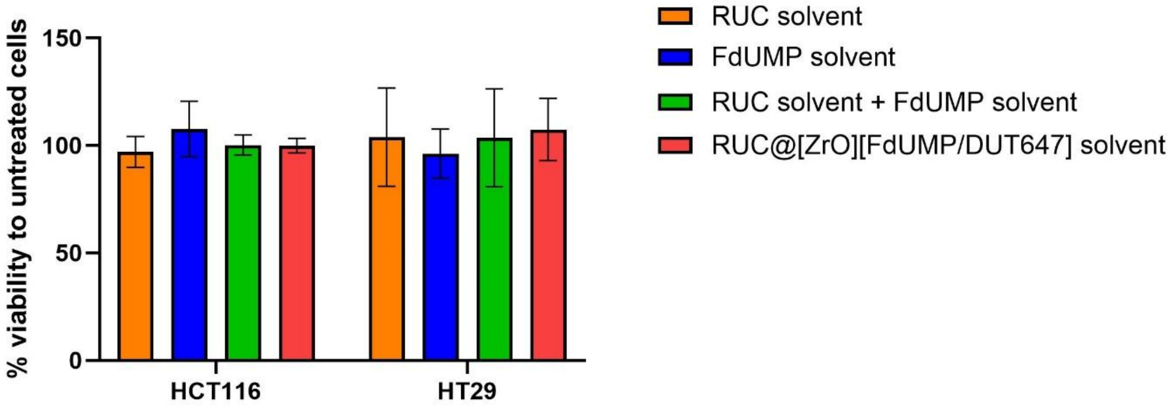


**Figure S16. Viability of HCT116 and HT29 cells following incubation with treatment solvents. Cell viability was measured after 72 h with an AlamarBlue HS assay.** Cells were incubated with the highest solvent concentrations used, which was 0.2 % DMSO for 100 µM RUC, 5 % water for 127 µM FdUMP, 0.2 % DMSO and 4 % water for 100 µM RUC and 100 µM FdUMP, and 773 µM trisodium citrate in 4.5 % water for RUC@[ZrO][FdUMP/DUT647] containing 100 µM RUC and 114 µM FdUMP. Values represent mean ± SD, and each experiment was carried out three times with three technical replicates.

To validate the observed tumor uptake of RUC@[ZrO][FdUMP/DUT647] in a HT29 xenograft model (*see main paper: Figure 6*), we performed the same staining and microscopic procedure on a C4-2 subcutaneous tumor of a SCID mouse without RUC@[ZrO][FdUMP/DUT647] injection (Figure S17). Bright RUC and DUT spots were clearly identifiable as artifacts (Fig. S17a) and unspecific DUT647 signal was detected outside of the tumor tissue (Figure S17b) However, no RUC or DUT647 signal was visible on tumor cells, confirming specific RUC@[ZrO][FdUMP/DUT647] uptake in Figure 7.

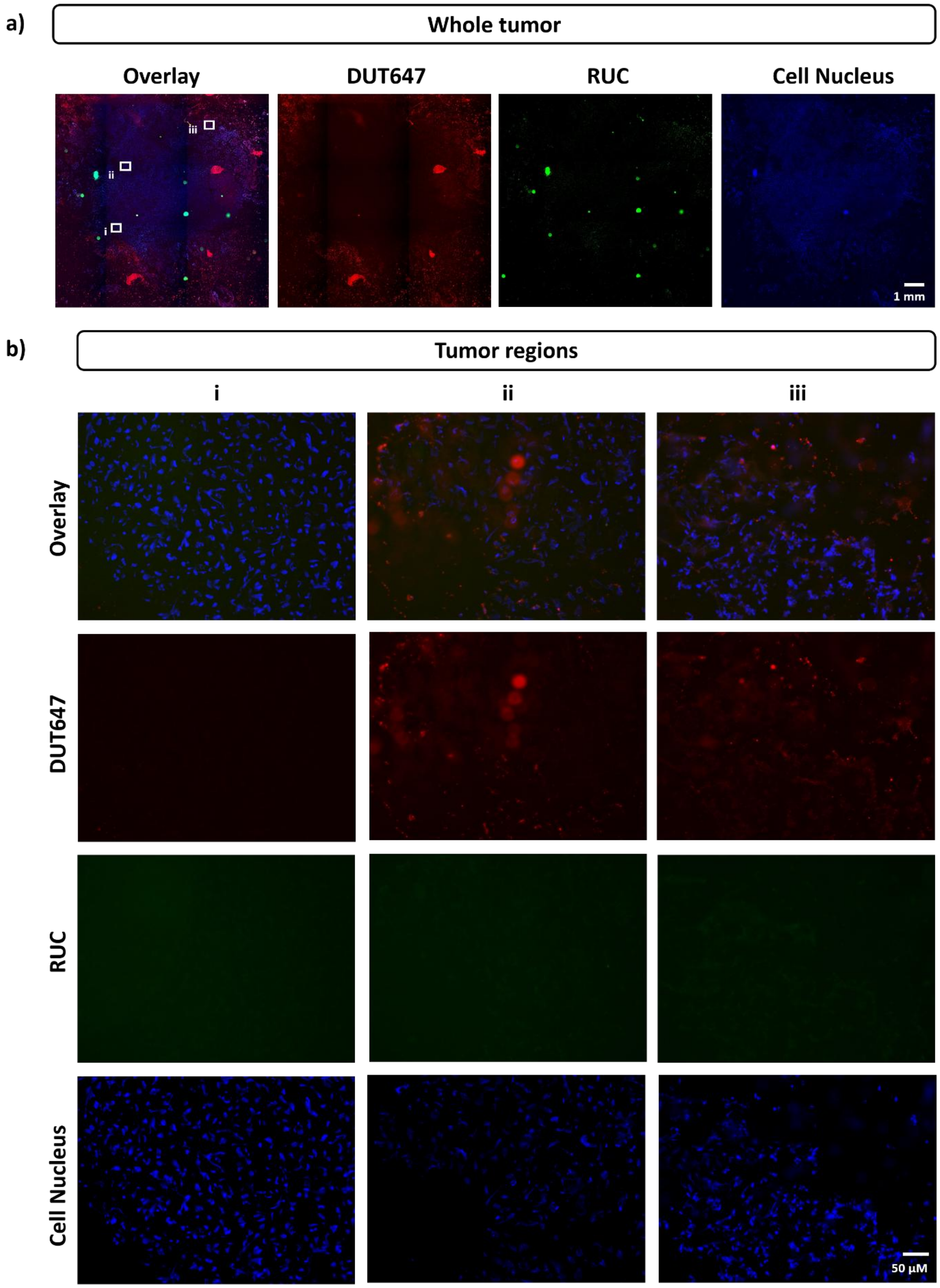


**Figure S17. Tumor microscopy images from one subcutaneous C4-2-tumor bearing SCID mouse without RUC@[ZrO][FdUMP/DUT647] injection**. The shell dye DUT647 of RUC@[ZrO][FdUMP/DUT647] (red), the core drug RUC of RUC@[ZrO][FdUMP/DUT647] (green) and cell nuclei (blue) are displayed. **a)** Overview image of the tumor showing no specific RUC or DUT647 uptake but bright punctuated artifacts. **b)** Three regions from a), indicated as i, ii, and iii, showing no RUC signal but DUT647 artifacts in areas outside the tumor mass.
